# Protein dynamics regulate distinct biochemical properties of cryptochromes in mammalian circadian rhythms

**DOI:** 10.1101/740464

**Authors:** Jennifer L. Fribourgh, Ashutosh Srivastava, Colby R. Sandate, Alicia K. Michael, Peter L. Hsu, Christin Rakers, Leslee T. Nguyen, Megan R. Torgrimson, Gian Carlo G. Parico, Sarvind Tripathi, Ning Zheng, Gabriel C. Lander, Tsuyoshi Hirota, Florence Tama, Carrie L. Partch

**Affiliations:** Department of Chemistry and Biochemistry, University of California Santa Cruz, Santa Cruz, CA 95064, USA; Institute of Transformative Bio-Molecules, Nagoya University, Nagoya 464-8601, Japan; The Scripps Research Institute, La Jolla, CA 92037, USA; Department of Pharmacology, University of Washington, Seattle, WA 98195, USA; Graduate School of Pharmaceutical Sciences, Kyoto University, Kyoto 606-8501, Japan; Howard Hughes Medical Institute, Box 357280, Seattle, WA 98125, USA; Department of Physics, Nagoya University, Nagoya 464-8601, Japan; RIKEN Center for Computational Science, Kobe 650-0047, Japan; Center for Circadian Biology, University of California San Diego, La Jolla, CA 92037, USA

**Keywords:** circadian rhythms, transcriptional regulation, cryptochrome, protein dynamics, Cryo-electron microscopy, protein crystallography, protein-protein interaction, molecular dynamics simulation

## Abstract

Circadian rhythms are generated by a transcription-based feedback loop where CLOCK:BMAL1 drive transcription of their repressors (PER1/2, CRY1/2), which bind to CLOCK:BMAL1 to close the feedback loop with ~24-hour periodicity. Here we identify a key biochemical and structural difference between CRY1 and CRY2 that underlies their differential strengths as transcriptional repressors. While both cryptochromes bind the BMAL1 transactivation domain with similar affinity to sequester it from coactivators, CRY1 is recruited with much higher affinity to the PAS domain core of CLOCK:BMAL1, allowing it to serve as a stronger repressor that lengthens circadian period. We identify a dynamic loop in the secondary pocket that regulates differential binding of cryptochromes to the PAS domain core. Notably, PER2 binding remodels this loop in CRY2 to enhance its affinity for CLOCK:BMAL1, explaining why CRY2 forms an obligate heterodimer with PER2, while CRY1 is capable of repressing CLOCK:BMAL1 both with and without PER2.

## Introduction

The circadian clock links our behavior and physiology to the daily light-dark cycle, providing a timekeeping system that ensures cellular processes are performed at an optimal time of day. At the cellular level, circadian rhythms are driven by a set of interlocked transcription-based feedback loops that take ~24 hours to complete. The basic helix-loop-helix PER-ARNT-SIM (bHLH-PAS) domain-containing transcription factor CLOCK:BMAL1 forms the positive arm of the core feedback loop (Gekakis et al., 1998). Several CLOCK:BMAL1 target genes (*Per1*, *Per2*, *Cry1*, *Cry2*) constitute the negative components that close the core feedback loop (Takahashi, 2017). PER and CRY proteins interact directly to form a large complex with the kinase CK1δ, and enter the nucleus after a delay to inhibit CLOCK:BMAL1 activity (Aryal et al., 2017; Lee et al., 2001). An additional feedback loop, comprising the nuclear receptors ROR and REV-ERB, controls the rhythmic expression of a subset of genes, including *Bmal1* (Preitner et al., 2002).

The two cryptochrome proteins, CRY1 and CRY2, appear to have distinct roles in the molecular circadian clock despite their high sequence and structural similarity (Michael et al., 2017b). Cryptochromes are essential for circadian rhythms, as *Cry1*^*−/−*^; Cry2^−/−^ double knockout mice are arrhythmic in constant darkness (van der Horst et al., 1999; Vitaterna et al., 1999). However, *Cry1*^*−/−*^ mice have a short period while *Cry2*^*−/−*^ mice have a long period, suggesting that they have non-redundant roles in the feedback loop. Early studies revealed that CRY1 is a stronger repressor of CLOCK:BMAL1 (Griffin et al., 1999). *Cry1* expression is also delayed with respect to *Cry2* and the *Per* genes (Lee et al., 2001; Ukai-Tadenuma et al., 2011), consistent with its recruitment to CLOCK:BMAL1-bound E-boxes in DNA in two distinct phases: a minor peak at circadian time (CT) 16-20 as part of the large PER-CRY repressive complexes (Aryal et al., 2017; Lee et al., 2001), and a major peak later at CT0-4 that is apparently independent of CRY2 and the PER proteins (Koike et al., 2012). However, the delayed timing of CRY1 expression does not exclusively account for its differential regulation of circadian period, because expressing *Cry2* from a minimal *Cry1* promoter in *Cry1*^*−/−*^; *Cry2*^*−/−*^; *Per2*^*Luc*^ fibroblasts or the suprachiasmatic nucleus (SCN) *ex vivo* still drives CRY2-like short periods (Edwards et al., 2016; Rosensweig et al., 2018). Several lines of evidence suggest that CRY1 has a biochemical activity not shared with CRY2. Fibroblasts, peripheral clocks and dissociated SCN neurons from *Cry1*^*−/−*^ *Per2*^*Luc*^ mice cannot generate sustained, cell-autonomous circadian rhythms (Liu et al., 2007) and the stabilization of CRY1, but not CRY2, in the SCN of *Fbxl3*^*Afh*^ mice selectively extends the duration of repression (Anand et al., 2013). Based on these collective observations, CRY1 may contribute to sustained cellular cycling through its ability to bind CLOCK:BMAL1 independently of PER proteins, helping to keep it in a poised and repressed state on DNA in the early morning (Koike et al., 2012) before the feedback loop begins anew.

What structural features of CRY1 and CRY2 might lead to their differential functions in the circadian clock? Both cryptochromes are defined by a highly conserved photolyase homology region (PHR) and divergent C-terminal tails that are intrinsically disordered (Partch et al., 2005). While deletion of the unstructured tails can modulate rhythm amplitude and period length, the PHR is required to generate circadian rhythmicity (Gao et al., 2013; Khan et al., 2012; Patke et al., 2017). We recently showed that CRY1 interacts directly with both CLOCK and BMAL1 through two distinct regions on the PHR (Michael et al., 2017a; Xu et al., 2015) (Figure 1A). The CC-helix of CRY1 and CRY2 are required for repression because it binds directly to the transactivation domain (TAD) of BMAL1 (Chaves et al., 2006; Czarna et al., 2011) (Figure 1B), sequestering the TAD from binding to coactivators (Xu et al., 2015). Repression also depends on the stable recruitment of CRY1 to the CLOCK:BMAL1 complex via the CLOCK PAS-B domain, which docks into the evolutionarily conserved secondary pocket on the CRY1 PHR (Michael et al., 2017a). Here we leverage these insights to identify the biochemical features that distinguish the different repressive capabilities of CRY1 and CRY2. We found that changes in the structure and dynamics of the serine loop, located adjacent to the secondary pocket, allow CRY1 to bind the CLOCK PAS-B domain within the PAS domain core of CLOCK:BMAL1 (Huang et al., 2012) with significantly higher affinity than CRY2. Moreover, modest substitutions in the serine loop and secondary pocket between CRY1 and CRY2 (Rosensweig et al., 2018) are sufficient to confer high affinity to the CRY2 PHR for the PAS domain core. Finally, we found that the CRY-binding domain (CBD) of PER2 has minimal effect on CRY1 affinity for CLOCK:BMAL1, but structurally remodels the serine loop in CRY2 to strengthen its binding to the CLOCK:BMAL1 PAS core. These data provide a biochemical rationale linking CRY2 repression to PER proteins, as well as explaining how CRY1 can act as a repressor of CLOCK:BMAL1 in the presence or absence of PER proteins.

**Figure 1.**
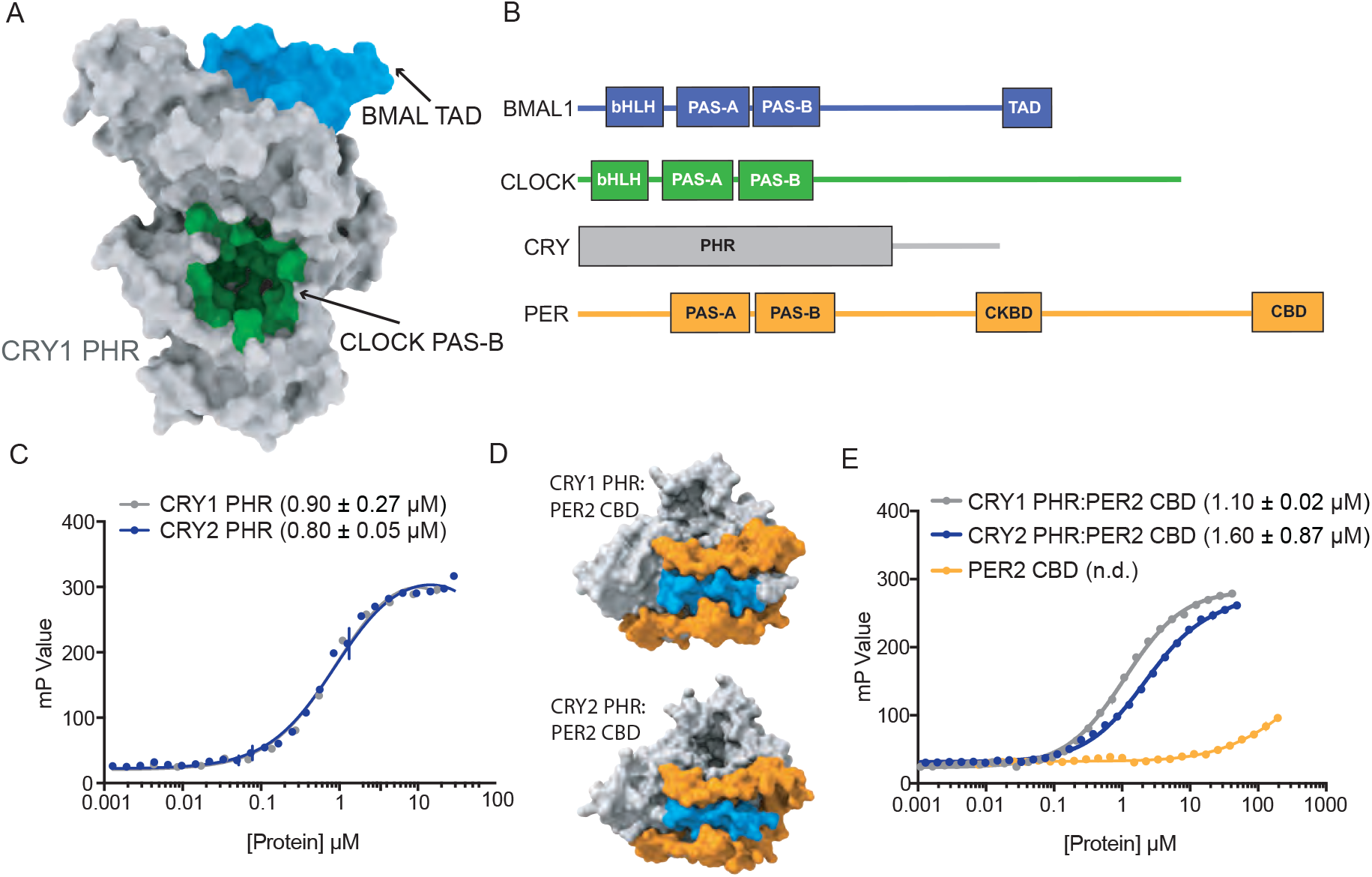
BMAL1 transactivation domain binding is comparable for the CRYPTOCHROME proteins. A, Crystal structure of CRY1 PHR (PDB: 5T5X) highlighting the BMAL TAD (blue) and CLOCK PAS-B (green) binding interface. B, Domain schematic of core clock proteins. bHLH, basic helix-loop-helix; PAS, PER-ARNT-SIM domain, TAD, transactivation domain; PHR, photolyase homology region; CKBD, Casein Kinase Binding Domain; CBD, CRY-binding domain. C, Fluorescence polarization (FP) assay of 5,6-TAMRA-labeled BMAL1 TAD (residues 594-626) binding to CRY1 (gray), replotted from (Gustafson et al., 2017), and CRY2 (blue). Mean ± SD data shown from one representative assay of n = 3 independent assays. Binding constants (mean ± SD) derived from n = 3 assays. D, Crystal structures of CRY1 PHR:PER2 CBD (PDB: 4CT0) and CRY2 PHR:PER2 CBD (PDB: 4U8H) with the PER2 CBD (orange) and CRY PHRs (gray), oriented to show the BMAL1 TAD binding interface (blue). E, FP assay of 5,6-TAMRA-labeled BMAL1 TAD binding to preformed CRY1 PHR:PER2 CBD (gray), CRY2 PHR:PER2 CBD (blue) or PER2 CBD (orange). Mean ± SD data shown from one representative assay of n = 3 independent assays. Binding constants (mean ± SD) derived from n = 3 assays. n.d., not determined.

## Results

### CRY1 and CRY2 bind similarly to the transactivation domain of BMAL1

We first compared the affinity of CRY1 and CRY2 for the BMAL1 TAD, as this represents a mechanism by which cryptochromes directly inhibit the activity of CLOCK:BMAL1 (Gustafson et al., 2017; Xu et al., 2015). Our fluorescence polarization-based binding assay with a TAMRA-labeled BMAL1 TAD probe showed that the CRY1 and CRY2 PHR domains bind with similar affinity (Figure 1C). However, the repressive clock protein complexes that accumulate early in the night contain CRYs with their associated PER proteins (Aryal et al., 2017), so it is important to understand how PER binding might influence CRY interactions with CLOCK:BMAL1. Crystal structures of the CRY1 and CRY2 PHR in their PER2-bound state show that the PER2 CBD wraps around the CC-helix of both CRYs (Figure 1D) (Nangle et al., 2014; Schmalen et al., 2014), coming into close proximity of the BMAL1 TAD binding site. To determine if PER2-bound CRY1 or CRY2 has different affinity for the BMAL1 TAD, we assessed binding to the TAMRA-BMAL1 TAD probe using CRY:PER2 CBD complexes as well as the PER2 CBD alone. Despite the intimate association of PER2 with the BMAL-binding site on CRY1 and CRY2, we found that both CRY:PER2 complexes bound the TAD similarly to the free CRYs, while the PER2 CBD displayed negligible affinity for the TAD by itself (Figure 1E). These data demonstrate that both CRY proteins have the capacity to sequester the TAD, either independently or in complex with the PER2 CBD, to repress CLOCK:BMAL1 activity.

### CRY1 binds substantially tighter than CRY2 to the PAS domain core of CLOCK:BMAL1

We next focused on the PAS domain core of CLOCK:BMAL1, as we previously showed that the PAS-B domain of CLOCK docks into an evolutionarily conserved secondary pocket on CRY1 (Michael et al., 2017a). To assess CRY binding, we initially used gel filtration chromatography with a CLOCK:BMAL1 construct that contains the structured basic helix-loop-helix DNA-binding domain and tandem PAS domains (bHLH-PAS) that form the ordered core of the heterodimer (Huang et al., 2012). Upon mixing the bHLH-PAS heterodimer with an excess of either CRY1 or CRY2 PHR, we observed that only CRY1 could bind CLOCK:BMAL1 tightly enough to co-migrate on the column (Figure 2A,B). These data suggested that there is a substantial difference in how the CRY PHRs interact with the PAS domain core of CLOCK:BMAL1. To quantitatively analyze this, we performed bio-layer interferometry (BLI) with the biotinylated tandem PAS-AB domain heterodimer, titrating in free CRY1 or CRY2 PHR to assess the kinetics of CRY binding (Figure 2C,D). We found that the CRY1 PHR binds with remarkably high affinity (*K*_d_ 65 ± 6 nM), while CRY2 bound with approximately 20-fold lower affinity (*K*_d_ 1.2 ± 0.2 μM).

**Figure 2.**
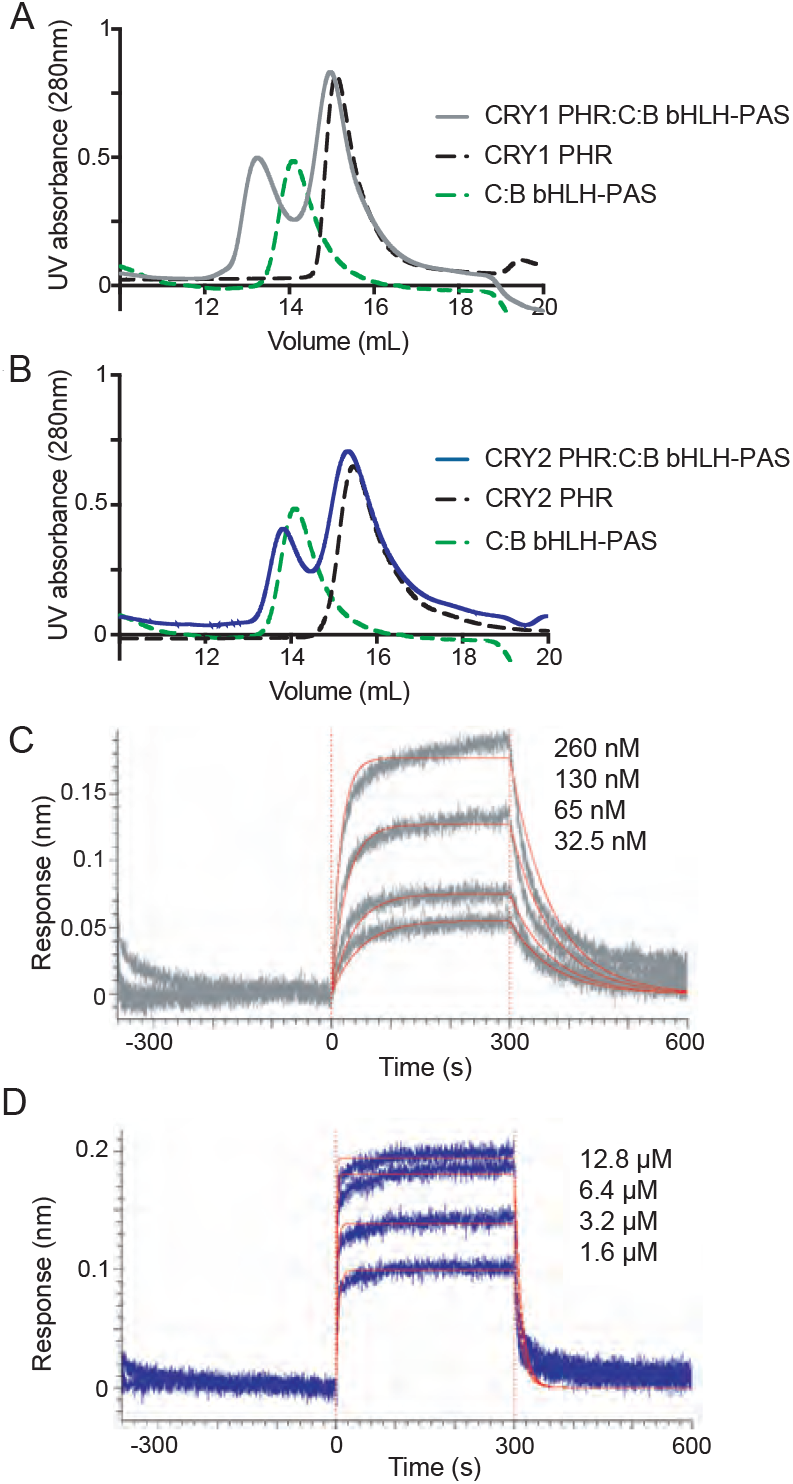
CRY1 binds 20 fold tighter to PAS domain core of CLOCK:BMAL1 than CRY2. A, Gel filtration analysis of complex formation of CRY1 PHR mixed with CLOCK:BMAL1 (C:B) bHLH-PAS. CRY1 PHR (black), C:B bHLH-PAS (green) and CRY1 PHR incubated with C:B bHLH-PAS (gray) were run on a S200 10/300 GL column. B, Gel filtration analysis of complex formation of CRY2 PHR mixed with CLOCK:BMAL1 (C:B) bHLH-PAS. CRY2 PHR (black), C:B bHLH-PAS (green) and CRY2 PHR incubated with C:B bHLH-PAS (blue) were run on a S200 10/300 GL column. C-D, BLI data for CRY1 PHR (gray, C) or CRY2 PHR (blue, D) binding to immobilized, biotinylated CLOCK:BMAL1 PAS-AB. Inset values represent the concentrations of CRY for individual binding reactions, top to bottom. Vertical red dashed lines indicate the beginning of association and dissociation. The red solid line is the nonlinear least squares fitting. CRY1 PHR *K*_*d*_ = 65 ± 6 nM; CRY2 PHR *K*_*d*_ = 1.2 ± 0.2 µM (mean of two independent experiments). Data shown from one representative experiment of n = 2 assays. See Figure S1 for additional information related to this figure.

The low micromolar affinity of CRY2 for the CLOCK:BMAL1 PAS domain core likely explains its inability to form a stable complex with CLOCK:BMAL1 under the conditions we used. We previously showed that multivalent interactions with both the PAS domain core and the BMAL1 TAD are important for transcriptional repression by CRY1 in cells (Xu et al., 2015). To see if reconstituting multivalent interactions by including the BMAL1 TAD could stabilize CRY2 binding to CLOCK:BMAL1, we explored complex formation in a sample containing CRY2 PHR with CLOCK bHLH-PAS and full-length BMAL1 by gel filtration. In contrast to the isolated bHLH-PAS core, the CRY2 PHR now co-eluted with the CLOCK:BMAL1 heterodimer, as visualized by SDS-PAGE of the peak fraction (Figure S1A). Therefore, although CRY2 binds with much lower affinity to the PAS core of CLOCK:BMAL1 than CRY1, the addition of a second binding interface with the BMAL1 TAD allows for stable complex formation with CRY2. Together, these data suggest that differential binding of CRY PHRs to the CLOCK:BMAL1 PAS domain core might underlie their different repressive roles in the circadian clock.

### The serine loop differentially gates access to the secondary pocket of CRY1 and CRY2

To explore the structural basis for differential binding of CRY PHRs to the CLOCK:BMAL1 PAS domain core, we examined existing crystal structures of mouse CRY1 and CRY2 PHR domains (Michael et al., 2017a; Xing et al., 2013) to identify sites of structural variability. While the overall structures are very similar (Cα Root Mean Square Deviation (RMSD) of 1.4 Å) (Figure 3A), molecular dynamics (MD) simulations suggested that CRY1 exhibits a higher degree of flexibility at the serine loop (Figure 3B) that is directly adjacent to the CLOCK-binding secondary pocket that we identified earlier (Michael et al., 2017a). Another loop (E196-S207) adjacent to serine loop shows lower flexibility in CRY2 simulations compared to CRY1. Proline residues at positions P219 and P225 in CRY2 replace an aspartate (D201) and serine (S207) in CRY1, which could be responsible for lower flexibility of this region in CRY2. The other side of the both CRYs also contained flexible regions, including the phosphate loop (p-loop) and a proximal loop containing a nuclear localization signal (NLS) (Figure 3B). In support of the dynamics observed in MD simulations, neither crystal structure of apo CRY1 contains density for the serine loop (Czarna et al., 2013; Michael et al., 2017a). However, this loop forms an α-helix in structures of apo or FAD-bound CRY2 (Xing et al., 2013). Only two amino acids differ between CRY1 and CRY2 in this loop: G43/A61 and N46/S64 (using CRY1/CRY2 numbering, Figure 3C). Since the substitution of alanine for glycine at position 61 likely stabilizes the helical content of CRY2 in this loop (Lopez-Llano et al., 2006), we created an *in silico* mutant of CRY2 swapping in the two CRY1 residues (A61G/S64N) to see if this would increase flexibility of the serine loop. Comparing the Root Mean Square Fluctuations (RMSF) for wild type and mutant CRY2 revealed that flexibility of the serine loop increased by maximum of 1 Å at A61G in the mutant (Figure 3D), suggesting that minor local perturbations to the serine loop could substantially impact CRY function. Reconstitution of *Cry1*^−/−^; *Cry2*^−/−^; *Per2*^Luc^ cells with a G43A/N46S CRY1 mutant (swapping CRY2 residues at this site into CRY1) lead to a significantly shorter period than wild-type CRY1 (Rosensweig et al., 2018), consistent with more CRY2-like weaker repressor activity (Hirota et al., 2012; Liu et al., 2007).

**Figure 3.**
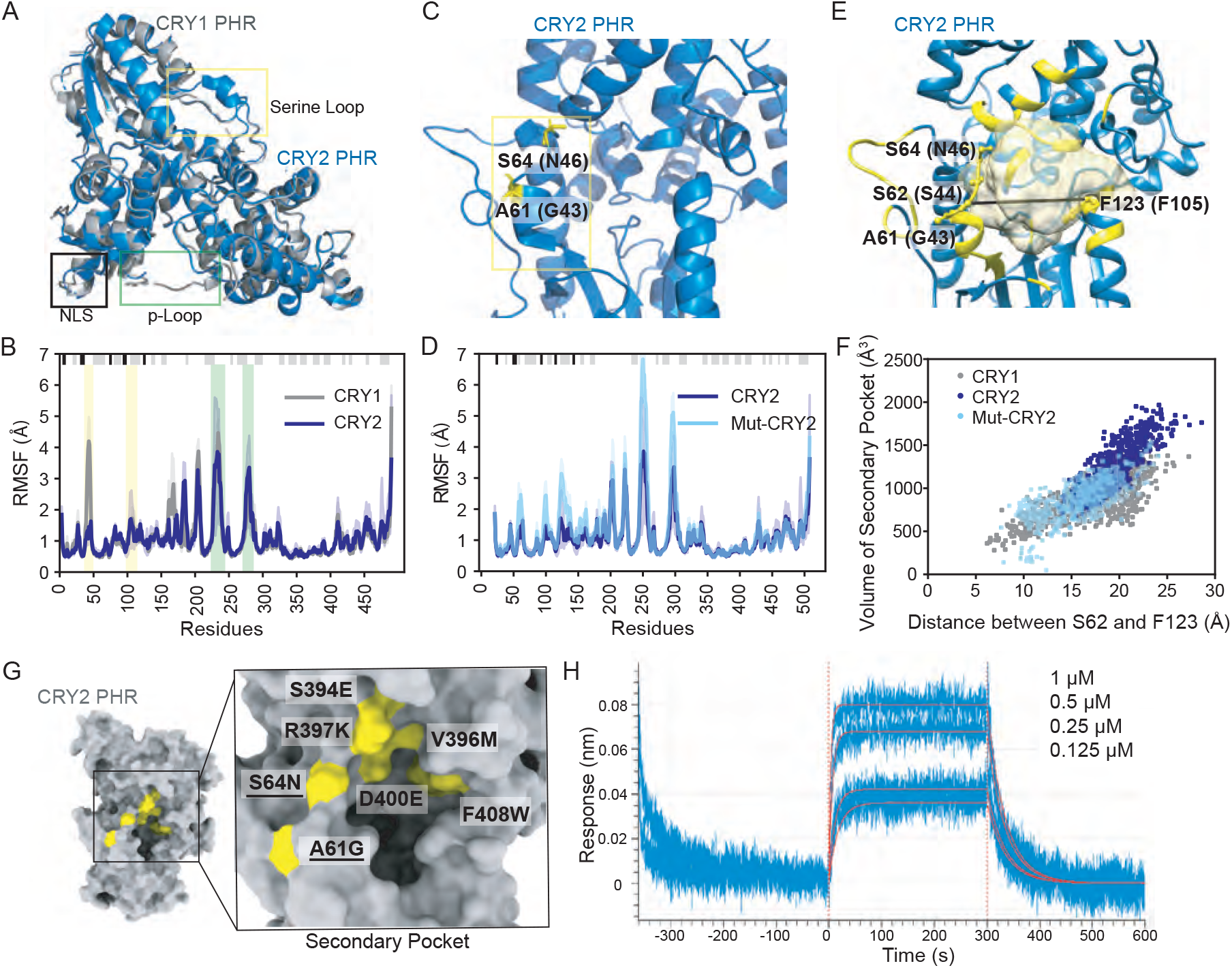
Flexibility of the serine loop controls CRY secondary pocket size and CLOCK:BMAL1 binding. A, Structural alignment of CRY1 PHR (PDB: 5T5X) and CRY2 PHR (PDB: 4I6E). Regions showing high structural variability have been boxed. B, Root mean square fluctuation (RMSF) values obtained from MD simulations for Cα atoms of each residue in CRY1 PHR (gray) or CRY2 PHR (blue) from n = 3 independent runs. The mean RMSF is depicted in dark shades with the variation between minimum and maximum values in light shades. Secondary structure is depicted at the top of the plot (α-helix, gray; β-strand, black) and residues in the serine loop and p-loop are shaded yellow and green, respectively. C, Crystal structure of CRY2 PHR (PDB: 4I6E) highlighting two residues that vary from CRY1 on the serine loop adjacent to the secondary pocket. D, RMSF values for the wild-type CRY2 PHR (dark blue) or Mut-CRY2 (S64N/A61G, light blue), as above. E, Volume of the secondary pocket (yellow cartoon and surface representation) and residues used for distance measurements of the secondary pocket opening in CRY2 (S62 and F123, depicted) or CRY1 (S44 and F105, by conservation). Black line indicates distance measured. F, Scatter plot of secondary pocket volume (Å^3^) versus opening distance between CRY2 S62 and F123 (or S44 and F105 in CRY1). G, Crystal structure of CRY2 PHR with the seven mutations (7M) identified in (Rosensweig et al., 2018). Underlined, the two mutations in the serine loop. H, BLI data for CRY2 7M (blue) binding to immobilized, biotinylated CLOCK:BMAL1 PAS-AB. Inset values represent the concentrations of CRY for individual binding reactions, top to bottom. Vertical red dashed lines indicate the beginning of association and dissociation. Red solid line, nonlinear least squares fitting to a one-site binding model. Calculated *K*_d_ for CRY2 7M PHR = 130 ± 12 nM (mean from n = 2 independent assays). See Figure S2 for additional information related to this figure.

To further explore the difference in the conformational ensemble of CRY1 and CRY2, we calculated the Kullback-Leibler (KL) divergence of the phi, psi, and chi1 dihedral angle distribution for all of the residues (Figure S2A). KL divergence is used to quantitatively describe the differences in the conformational states of residues in two equilibrium ensembles (McClendon et al., 2012). A high KL divergence for a particular residue signifies that its conformational state in torsional space shows considerable difference between the two ensembles. Several residues throughout CRY1 and CRY2 show high KL divergence; in particular, the residues with the highest divergence concentrate in and around the secondary pocket, corresponding to CRY1 residues W52, Q79, D82, Y100, R127, W390 (Figure S2A). Notably, the comparison of conformational ensembles for wild-type CRY2 and the A61G/N64S CRY1-swapped CRY2 mutant also reveal high divergence at the secondary pocket and, to a lesser extent, throughout the rest of the protein (Figure S2B). These data suggest that the effect of substitutions in the secondary pocket that alter its rigidity may extend beyond the secondary pocket to influence the overall dynamical ensemble of CRY PHRs.

Based on these data, alterations in the structure and dynamics of the serine loop likely influence access to the secondary pocket that is located immediately adjacent to this loop. To quantify how variations in the serine loop of CRY1/CRY2 alter the conformational landscape of the secondary pocket, we calculated the volume of the pocket and the distance between center of mass of residue S62 and F123 in CRY2 (corresponding to S44 and F105 in CRY1), located on the opposite sides of the pocket, for each of the independent MD trajectories (Figure 3E). This analysis revealed an approximately linear relationship between the interatomic distance of the loops that line the secondary pocket and its volume (Figure 3F). The overall distribution for CRY2 measurements clustered toward the upper right region of this plot, demonstrating that CRY2 samples a larger pocket volume (1215 ± 275 Å^3^) as well as having larger opening to the pocket. The distribution of CRY1 measurements revealed a larger variation in distance across the pocket, likely due to the flexibility of the serine loop, but consistently had a smaller pocket volume (876 ± 217 Å^3^, p<0.001, Wilcoxon ranksum test) compared to CRY2. As predicted from its enhanced flexibility, the *in silico* mutant of the CRY2 serine loop (i.e., A61G/N64S) resulted in a significant change in its overall distribution towards a smaller volume of the secondary pocket (868 ± 230 Å^3^, p<0.001, Wilcoxon ranksum test) and shorter distances across the pocket, much like we found for CRY1. These data suggest that decreasing the flexibility of the serine loop correlates with increased volume of the secondary pocket and might play a role in the decreased affinity of CRY2 for the PAS domain core of CLOCK:BMAL1.

We sought to experimentally test if mutations on the serine loop and around the secondary pocket altered the ability of CRY2 to bind the PAS domain core of CLOCK:BMAL1. We utilized the CRY2 7M mutant (Rosensweig et al., 2018), which incorporates seven mutations in and around the secondary pocket that swap CRY2 residues with corresponding residues on CRY1 (Figure 3G). Genetic reconstitution of circadian rhythms in the CRY2 7M mutant (under control of the minimal *Cry1* promoter (Ukai-Tadenuma et al., 2011)) in *Cry1*^*−/−*^; *Cry2*^*−/−*^; *Per2*^*Luc*^ cells demonstrated that CRY2 7M was sufficient to largely recapitulate a CRY1-like repression phenotype and period (Rosensweig et al., 2018), suggesting that a change in the secondary pocket is sufficient to account for most differences in activity of the two CRY isoforms in the molecular clock. In line with this, we found that the CRY2 7M mutant had a ten-fold enhancement in affinity for the CLOCK:BMAL PAS domain core, as determined by BLI (Figure 3H). These data demonstrate that modest sequence differences in the secondary pocket of CRY1 and CRY2 can lead to significant differences in their ability to bind CLOCK:BMAL1 and could therefore influence their roles as circadian repressors in the mammalian circadian clock.

### The CRY1 serine loop is disordered in the CRY1 PHR:PER2 CBD complex

The PER2 CBD has an intimate association with the BMAL1 TAD-binding site at the C-terminal CC helix of CRY1 and CRY2 (Figure 1D), but it also wraps around the distal side of the PHR to come into close contact with the secondary pocket (Nangle et al., 2014; Schmalen et al., 2014) (Figure 4A). Because of its proximity to the secondary pocket, we wanted to see if PER2 might play a role in regulating the structure of the serine loop to control the affinity of CRY1 or CRY2 for CLOCK:BMAL1. However, a previous crystal structure of the CRY1 PHR:PER2 CBD complex (PDB: 4CT0) contained a non-native sequence that packed against the secondary pocket, resulting in non-physiological ordering of the dynamic serine loop in CRY1 (Schmalen et al., 2014) (Figure 4B). In order to see if the PER2 CBD alters the serine loop and secondary pocket of CRY1, we determined a crystal structure of the CRY1 PHR:PER2 CBD complex that eliminates this artifact (Figure 4A, Figure S3). The overall conformation of the PER2 CBD in our structure is highly similar to the previously solved structure (0.76 Å Cα RMSD over residues 1136-1207). Although the PER2 CBD passes near the secondary pocket of CRY1, it makes only one direct contact with the serine loop where the backbone of PER2 Gln1135 forms a hydrogen bond with Ser45 on the loop (Figure 4C, Figure S6A). As a result, we observed modest ordering of the serine loop in CRY1, suggesting that it maintains some of its flexibility in the PER2-bound state.

**Figure 4.**
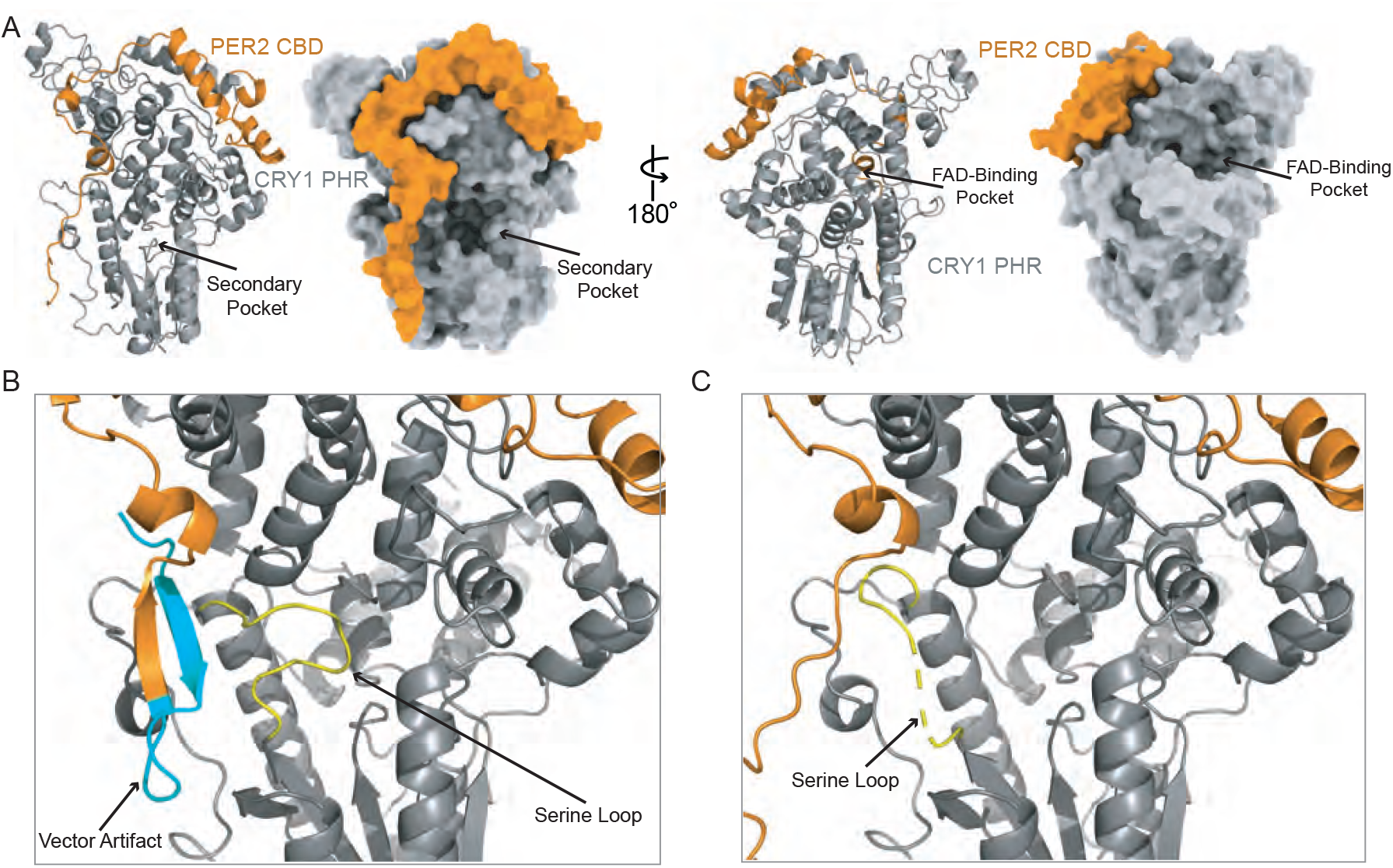
CRY1 PHR:PER2 CBD crystal structure. A, Crystal structure of the CRY1 PHR:PER2 CBD complex (PDB: 6OF7) with CRY1 PHR (gray) and PER2 CBD (orange). Ribbon and surface representations on the left highlight the secondary pocket, while the FAD binding pocket is shown on the right. B, Zoomed in view of the secondary pocket of the previously published CRY1 PHR:PER2 CBD crystal structure (PDB: 4CT0) (Schmalen et al., 2014) with the serine loop (yellow) that is ordered by contact with the vector artifact (cyan). C, Same view of the secondary pocket in our crystal structure (PDB: 6OF7) showing the partially ordered serine loop (yellow). See Figure S3 and supplemental table 1 for additional information related to this figure.

### Visualizing the architecture of a PER2 CBD:CRY1 PHR:CLOCK PAS-B complex

We previously used small-angle x-ray scattering (SAXS) studies of the CRY1 PHR with the CLOCK:BMAL1 bHLH-PAS heterodimer, along with site-directed mutagenesis of the predicted CRY1 PHR:CLOCK PAS-B interface, to model how CRY1 binds to the PAS domain core of CLOCK:BMAL1 (Michael et al., 2017a). However, some ambiguities in our computationally-derived HADDOCK model of the CRY1 PHR:CLOCK PAS-B complex and the low-resolution of SAXS data prevented us from confidently identifying a preferred model of the ternary complex from several possibilities that were consistent with the scattering data. We attempted to resolve this ambiguity through single particle cryo-electron microscopy (cryo-EM) analyses of the complex formed between the CLOCK:BMAL1 bHLH-PAS heterodimer bound to a short DNA duplex containing the *Per2* E-box, and the circadian repressors CRY1 PHR:PER2 CBD.

Although the inherent conformational heterogeneity of the complex prevented determination of a high-resolution structure, 2D averages and a low-resolution 3D reconstruction provide important architectural insights (Figure 5A and Figure S4). Cryo-EM analyses demonstrate that the complex consists of a smaller globular volume protruding from the concave side of an elongated volume. The molecular envelope of the low-resolution cryo-EM reconstruction was compared to existing crystal structures, and we concluded that the elongated volume likely corresponds to the CRY1 PHR:PER2 CBD complex and that the smaller globular region is likely an interacting PAS domain from either CLOCK or BMAL1. Our previous biochemical data and integrative HADDOCK modeling (Michael et al., 2017a) indicated that the PAS-B domain from CLOCK, not BMAL1, interfaces with the CRY1 PHR, and rigid body docking of crystal structures into our reconstruction support this arrangement. A comparison of 2D projections of our rigid body-docked model (see Methods) with the 2D averages obtained from our cryo-EM data further validated our proposed architecture (Figure S4E). We were unable to identify densities that correspond to BMAL1, the bHLH and PAS-A domains of CLOCK, or the E-box DNA, likely due to the high degree of internal flexibility of the CLOCK:BMAL1 heterodimer, which we previously observed in SAXS studies (Michael et al., 2017a). We also observed that the PAS-B domains of CLOCK and BMAL1 undock from one another upon binding to CRY1 (Michael et al., 2017a), which is consistent with our identification of a single PAS-B domain in our reconstruction.

**Figure 5.**
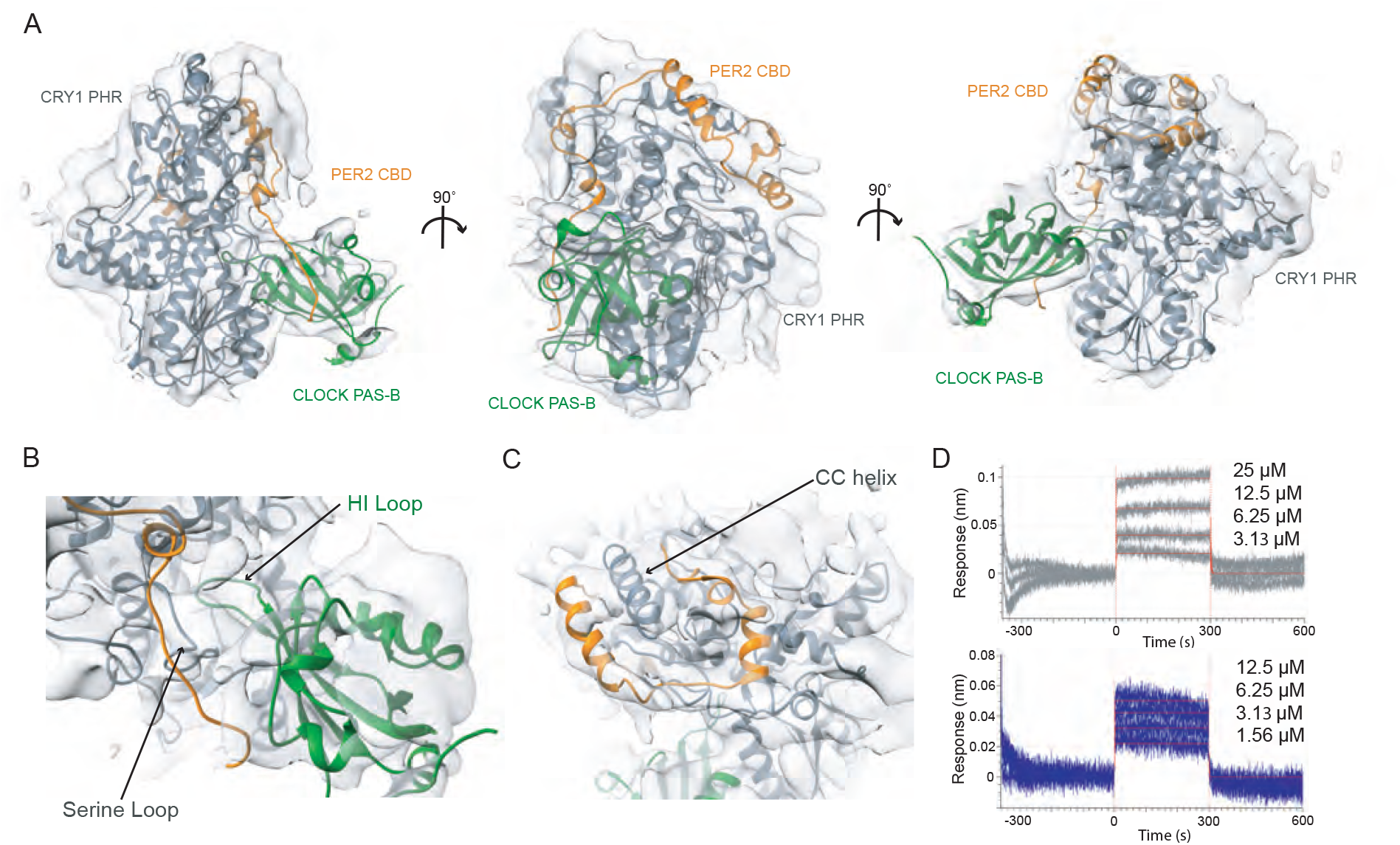
Cryo-EM structure of PER2 CBD:CRY1 PHR:CLOCK PAS-B. A, Cryo-EM map of the PER2 CBD:CRY1 PHR:CLOCK PAS-B complex with EM density rendered semi-transparent with the pseudoatomic model of CRY1 PHR (gray), PER2 CBD (orange), and CLOCK PAS-B (green) docked in. B, Zoomed in view of the region containing the serine loop near the secondary pocket where CLOCK PAS-B docks into CRY1. C, Zoomed in view of the PER2 CBD wrapping around the CC-helix of CRY1. D, BLI data for CRY1 PHR (gray) or CRY2 PHR (blue) binding to immobilized, biotinylated W362A CLOCK:BMAL1 PAS-AB. Inset values represent the concentrations of CRY for individual binding reactions, top to bottom. Vertical red dashed lines indicate the beginning of association and dissociation. Red solid line, nonlinear least squares fitting to a one-site binding model. Calculated *K*_d_ for CRY1 PHR = 6.6 ± 2.6 μM, CRY2 PHR = 10.2 ± 0.2 μM (mean from n = 2 independent assays). See Figures S4, S5 and supplemental table 2 for additional information related to this figure.

Given the challenges we faced with interdomain flexibility of the PER2 CBD:CRY1 PHR: CLOCK-BMAL1 bHLH-PAS complex, we used an integrative structural biology approach to model the portion of the complex included in our experimental data. Pseudoatomic HADDOCK models of the CRY1 PHR:CLOCK PAS-B were obtained using CRY1 representative structures from MD simulations. We superimposed representative structures from the three largest clusters obtained from HADDOCK onto the CRY1 PHR:PER2 CBD crystal structure. Among the representative structures of top three clusters, Cluster 3 had significant steric hindrance to the bound PER2 CBD (Figure S5). Consequently, only structures from the top two largest clusters, showing binding modes with the least steric hindrance between PER2 CBD and CLOCK PAS-B, were docked into the cryo-EM density map via rigid body fitting (Figure S5). A representative structure from the largest cluster with the lowest restraint violation (Cluster 1) provided the best fit (Figure S5). This model was further improved by performing flexible fitting using MDFit (Whitford et al., 2011), which modified the conformation of the CRY1 PHR:PER2 CBD:CLOCK PAS-B complex to fit the EM map (Figure 5A-C).

A close-up view of the CRY1 secondary pocket in the EM model shows density for the PER2 CBD and/or the serine loop where the PER2 CBD, CRY1, and CLOCK PAS-B are in close proximity (Figure 5B). The structural model provides experimental evidence that the HI loop of CLOCK PAS-B docks into the CRY1 secondary pocket, which was predicted by our earlier computational modeling and biochemical analyses (Michael et al., 2017a). We previously showed that mutation of W362 to alanine reduced binding of the CRY1 PHR to the isolated PAS-B of CLOCK or the CLOCK:BMAL1 PAS domain core and significantly decreased CRY1 repression of CLOCK:BMAL1 in 293T cells (Michael et al., 2017a). To examine how this CLOCK mutation quantitatively influences binding to CRY1 and CRY2, we performed BLI-based binding studies using a PAS domain core harboring the W362A CLOCK mutant (Figure 5D). Both CRY1 and CRY2 PHR demonstrated significantly reduced affinity for the mutant PAS domain core, with a *K*_d_ for CRY1 of 6.6 ± 2.6 μM, down nearly 100-fold from the native affinity, while CRY2 bound with 10.2 ± 0.2 μM affinity, 9-fold reduction compared to wild-type affinity. These results demonstrate the critical role of CLOCK PAS-B in the interaction with both CRY1 and CRY2 with the CLOCK:BMAL1 PAS domain core, specifically highlighting the critical role of the tryptophan residue on the HI loop of CLOCK PAS-B.

### The PER2 CBD remodels the CRY2 serine loop to promote binding to the PAS domain core of CLOCK:BMAL1

Prior observations suggest that CRY2 may have a need to associate with PER proteins to form a functional repressive complex, while CRY1 can bind to CLOCK:BMAL1 independent of the PER proteins (Koike et al., 2012; Rosensweig et al., 2018). To address this biochemical model, we explored how PER2 binding might regulate affinity of the CRY1 and CRY2 PHR for the PAS domain core of CLOCK:BMAL1. Comparing the secondary binding pocket of apo CRY1 PHR structure to our CRY1 PHR:PER2 CBD structure (Figure 6A), we observed that PER2 CBD induces a modest structuring at the very end of the loop (residues 44-47). This reorganization is caused by an interaction between Ser45 on CRY1 and the backbone of Gln1135 on the PER2 CBD (Figure S6A). To test how the addition of PER2 CBD alters the binding between CRY1 and CLOCK:BMAL1 PAS-AB, we utilized BLI binding assays using preformed CRY:PER2 CBD complexes (Figure 6B, S6B) with the immobilized PAS domain core. We found that CRY1:PER2 CBD binds to CLOCK:BMAL1 with about 3-fold decreased affinity. Therefore, addition of the PER2 CBD weakens the interaction of the CRY1 PHR with CLOCK:BMAL1 PAS core, indicating that this modest ordering of the CRY1 serine loop may make the interaction with CLOCK PAS-B less favorable.

**Figure 6.**
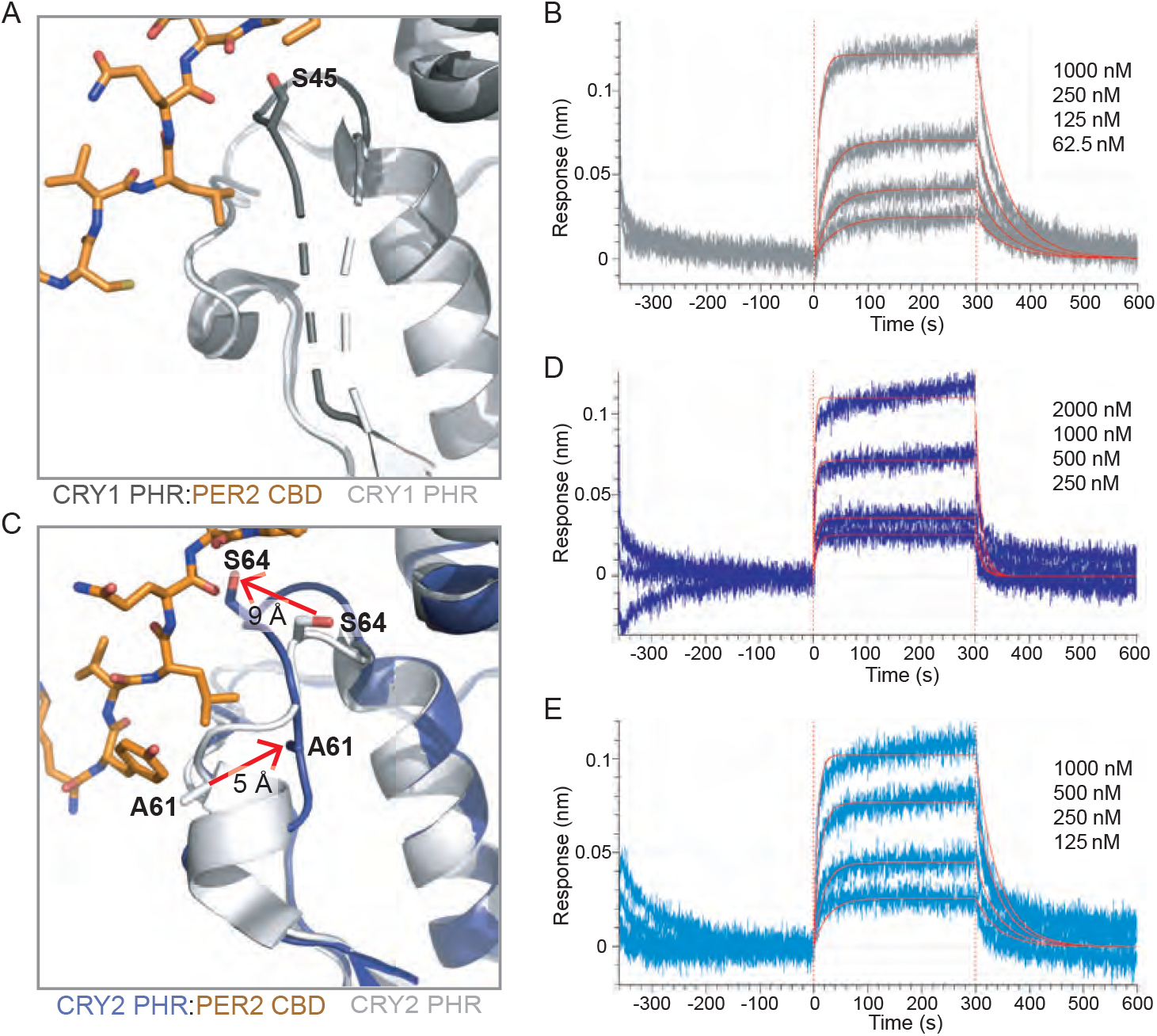
PER2 CBD alters the serine loops of CRY1 and CRY2 altering their affinity for the CLOCK:BMAL1 PAS core. A, Comparison of the serine loop from apo CRY1 PHR (light gray, PDB: 5T5X) and the CRY1 PHR:PER2 CBD complex (dark gray:orange, PDB: 6OF7). A slight structuring of the C-terminus of the serine loop occurs upon addition of PER2 CBD, but the N-terminus remains flexible (dashed line). B, BLI data for the CRY1 PHR:PER2 CBD complex (gray) binding to immobilized, biotinylated CLOCK:BMAL1 PAS-AB. Inset values represent the concentrations of CRY for individual binding reactions, top to bottom. Vertical red dashed lines indicate the beginning of association and dissociation. Red solid line, nonlinear least squares fitting to a one-site binding model. Calculated *K*_d_ for CRY1 PHR:PER2 CBD = 196 ± 34 nM (mean from n = 2 independent assays). C, Comparison of the serine loop from the apo CRY2 PHR (light gray, PDB: 4I6E) and the CRY2 PHR:PER2 CBD complex (blue:orange, PDB: 4U8H). Addition of the PER2 CBD alters the structure of the serine loop, decreasing the helical content and pulling residue S64 ~9Å away from the secondary pocket, and pushing A61 ~5 Å into the pocket. D, BLI data for the CRY2 PHR:PER2 CBD complex (dark blue) binding to immobilized, biotinylated CLOCK:BMAL1 PAS-AB domains. Calculated *K*_d_ for CRY2 PHR:PER2 CBD = 604 ± 29 nM (mean from n = 2 independent assays). E, BLI data for the CRY2 PHR 7M:PER2 CBD complex (light blue) binding to immobilized, biotinylated CLOCK:BMAL1 PAS-AB. Calculated *K*_d_ for CRY2 PHR 7M:PER2 CBD = 159 ± 66 nM (mean from n = 2 independent assays). See Figure S6 for additional information related to this figure.

By contrast, our comparison of the apo CRY2 PHR structure with the CRY2 PHR:PER2 CBD complex revealed a major structural reorganization of the serine loop by PER2 (Figure 6C). The serine loop in apo CRY2 PHR is structured with a short α-helix in the N-terminus of the serine loop and a stable, but extended loop at the C-terminus (Nangle et al., 2014; Xing et al., 2013). In the CRY2 PHR: PER2 CBD complex, the C-terminus of the serine loop is stabilized in an alternate position by multiple hydrogen bonds between Ser64 on CRY2 and Gln1135/Asp1136 of PER2 CBD (Figure S6C). This interaction moves Ser64 of CRY2 out of the pocket by approximately 9 Å (Figure 6C) and largely mimics the PER2-dependent interaction and subsequent loop orientation we observed with Ser45 on CRY1 (Figure S6A). Additionally, the PER2 CBD remodels the N-terminus of the serine loop, causing an unfolding of the short α-helix and a translation of Ala61 ~5 Å towards the secondary pocket (Figure 6C). Therefore, the addition of PER2 CBD causes the serine loops of CRY1 and CRY2 to take on a similar conformation, with the loop in CRY1 becoming more structured and the loop in CRY2 becoming less structured. We then utilized BLI binding assays to quantitatively assess how addition of the PER2 CBD and its subsequent structural rearrangements would alter CRY2 affinity for the immobilized PAS domain core of CLOCK:BMAL1. We found that formation of a stable CRY2:PER2 CBD complex enhanced its affinity for CLOCK:BMAL1 2-fold (Figure 6D), consistent with previous data showing that addition of PER2 could enhance co-immunoprecipitation of CLOCK:BMAL1 by CRY2 (Rosensweig et al., 2018). To confirm that the effect of PER2 CBD is due to regulation of the CRY serine loop and not a direct interaction with CLOCK:BMAL1, we also probed for a direct interaction of PER2 CBD with the PAS domain core of CLOCK:BMAL1 by BLI and observed no detectable binding (Figure S6D). These results demonstrate that addition of PER2 makes CRY2 a stronger repressor by enhancing its affinity for CLOCK:BMAL1, although it still exhibits weaker binding than CRY1, either with or without the PER2 CBD.

Based on our structural analyses, we attribute this tighter CLOCK:BMAL1 binding to the loss of helical structure in the serine loop observed when CRY2 is bound to the PER2 CBD. Therefore, the CRY2 7M mutant, which substitutes CRY1 residues into the serine loop and secondary pocket, might be expected to not exhibit a gain in affinity for CLOCK:BMAL1 in the presence of the PER2 CBD. To test this prediction, we collected BLI binding data on the CRY2 7M mutant in the presence of PER2 CBD. In line with this model, the CRY2 7M:PER2 CBD complex did not exhibit a gain in affinity for CLOCK:BMAL1 (Figure 3E). These data demonstrate that PER2 CBD tunes affinity of the CRY PHRs for the PAS domain core of CLOCK:BMAL1 through differential interactions at the serine loop.

Notably, when CRY1 or CRY2 are associated with their co-repressor PER2, their affinity for CLOCK:BMAL1 is at its most similar (within a ~3-fold difference) compared to the 20-fold difference that we observe with the two proteins in the absence of PER2. This provides biochemical justification supporting the *in vivo* observation that CRY1 can bind CLOCK:BMAL1 both with and without PER proteins, whereas CRY2 may require PER1 or PER2 to form a functional repressive complex (Koike et al., 2012; Rosensweig et al., 2018).

## Discussion

Since the identification of mammalian core clock proteins approximately two decades ago, numerous studies have helped to piece together how they work at the network level to generate the feedback loop that confers circadian timekeeping (Takahashi, 2017). However, we still lack a mechanistic understanding of how most clock proteins modify and/or interact with one another to fulfill their important roles as dedicated cogs in the molecular clock. In this study, we used biochemistry, structural biology, and computational methods to explain how CRY1 and CRY2 association with CLOCK:BMAL1 is regulated by the serine loop. In contrast to CRY2, CRY1 has a dynamic serine loop that confers tight binding to CLOCK at the secondary pocket. Remarkably, we found that association with PER2 unfolds and remodels the ordered serine loop of CRY2 to enhance its affinity for CLOCK. We confirmed that the serine loop and secondary pocket are sufficient to regulate association with the PAS domain core of CLOCK:BMAL1 using the CRY2 7M mutant (Rosensweig et al., 2018), which confers CRY1-like affinity and dynamics to the serine loop, and importantly, resistance to the PER2-mediated enhancement of affinity that we observed with CRY2. Therefore, small changes in the structure and dynamics of the serine loop with and without PER2 allow cryptochromes to fine-tune their repressive power by controlling how tightly they bind to CLOCK:BMAL1.

Our data suggest that the differential effect of the PER2 CBD on CRY PHR binding to CLOCK:BMAL1 allows it to serve as a molecular equalizer— the PER2 CBD compensates for poorer binding by CRY2 by modestly enhancing its affinity for the transcription factor, while also modestly reducing CRY1 affinity to bring the overall affinities of the two cryptochromes for the PAS domain core of CLOCK:BMAL1 to similar levels. This suggests that the actions of CRY1 and CRY2 within the heteromultimeric PER-CRY repressive complexes (Aryal et al., 2017; Lee et al., 2001) may be largely similar. The apparent dependency of CRY2 on PER1/2 (the PER isoforms that possess a CRY-binding domain (Miyazaki et al., 2001; Yagita et al., 2002)) is consistent with its expression pattern, which occurs in phase with the PER proteins, whereas the peak expression of CRY1 is delayed by several hours (Lee et al., 2001; Ukai-Tadenuma et al., 2011). The same temporal profile is observed in their genome-wide mapping of DNA occupancy with CLOCK:BMAL1, revealing two temporally distinct waves; ‘early’ repressive complexes that contain PER proteins, their associated kinase CK1δ, CRY1, CRY2, and variable association of additional epigenetic regulators (Aryal et al., 2017; Kim et al., 2015), and a ‘late’ repressive complex of CRY1 bound to CLOCK:BMAL1 in the apparent absence of PER proteins (Koike et al., 2012; Partch et al., 2014). The 20-fold increase in affinity that we observed with CRY1 for the PAS domain core of CLOCK:BMAL1 relative to CRY2 likely plays a critical role in its ability to participate in the ‘late’ complex, and may also give rise to the CRY1-selective extension of the repression phase observed in the *Fbxl3*^*Afh*^ mutant (Anand et al., 2013). Altogether, these data suggest that a proper balance of the biochemically distinct early and late repressive complexes work together to generate of the circadian period of ~24 hours.

Understanding precisely why the more dynamic serine loop helps CRY1 bind to the PAS domain core of CLOCK:BMAL1 awaits determination of a higher resolution structure. Like many other transcription factors, CLOCK:BMAL1 is a ‘malleable machine’ and its dynamic nature presents challenges for structure determination (Fuxreiter et al., 2008). From the perspective of cryptochromes, the dynamic nature of the serine loop may be associated with evolution of transcriptional repressor function. Notably, the secondary pocket in photoreceptive cryptochromes from *Arabidopsis* and Drosophila (Brautigam et al., 2004; Levy et al., 2013), as well as the evolutionarily related DNA repair enzymes, CPD or (6-4) photolyase (Hitomi et al., 2009; Park et al., 1995), is considerably smaller in volume, where it binds to small molecule cofactors that facilitate light harvesting (Sancar, 2003). The ability of repressor-type cryptochromes to interact with CLOCK PAS-B appears to be deeply rooted in evolution of the metazoan circadian clock, because the vertebrate-like cryptochromes that act as transcriptional repressors in insects (Zhu et al., 2005) like the monarch butterfly also depend to the same extent on multivalent interactions with CLOCK PAS-B and the BMAL1 TAD as they do in mammals (Sato et al., 2006; Xu et al., 2015; Zhang et al., 2017).

The BMAL1 TAD is the primary driver of transcriptional activation by CLOCK:BMAL1 in mammalian and insect systems with repressor-type cryptochromes; truncation or mutation of the TAD decimates CLOCK:BMAL1 activity and leads to arrhythmicity *in vivo* (Kiyohara et al., 2006; Park et al., 2015; Zhang et al., 2017). Sequestration of the TAD by repressor-type cryptochromes allows them to compete with coactivators at a highly conserved and overlapping binding motif (Xu et al., 2015). Strikingly, we did not observe differential binding for the BMAL1 TAD by CRY1 or CRY2 in the absence or presence of the PER2 CBD. This suggests that the changes in CRY affinity for the PAS domain core of CLOCK:BMAL1 that we observed underlie the major functional differences between CRY1 and CRY2 that contribute to differential period regulation (Edwards et al., 2016; van der Horst et al., 1999; Vitaterna et al., 1999) and the ability of CRY1 to sustain cellular circadian rhythms (Liu et al., 2007). Therefore, cryptochrome recruitment to the PAS domain core likely represents a key regulatory node in the clock and may play a role in an inherited *Cry1* variant that modulates affinity for CLOCK:BMAL1 to alter circadian timing in humans (Patke et al., 2017).

## Acknowledgements

We would like to thank the beamline staff at 23-ID-D for their assistance during data collection at the Advanced Photon Source. We thank J.C. Ducom at Scripps Research High Performance Computing and C. Bowman at Scripps Research for computational support, as well as B. Anderson at the Scripps Research Electron Microscopy Facility for microscopy support. This work was funded by National Institutes of Health Grant R01 GM107069 (to C.L.P.), DP2 EB020402 (to G.C.L), RIKEN Dynamic Structural Biology Project (to F.T.). G.C.L. is supported by the Pew Charitable Trusts as a Pew Scholar and an Amgen Young Investigator award. J.L.F. was supported by the UC Office of the President and UCSC Chancellor’s Postdoctoral Fellowship. C.R.S. was supported by a National Science Foundation predoctoral fellowship. Computational analyses of EM data were performed using shared instrumentation funded by NIH S10OD021634 to G.C.L.

## Author contributions

Conceptualization, J.L.F., A.K.M., A.S., T.H., F.T. and C.L.P.; Investigation, J.L.F., A.S., C.R.S., A.K.M., P.L.H., S.T., C.R., and M.T.; Resources, L.T.N. and G.C.P.; Validation, J.L.F., A.S., and C.R.S.; Writing – Original Draft, J.L.F., A.S., C.R.S., and C.L.P.; Writing – Review & Editing, J.L.F., A.S., C.R.S., P.L.H., G.C.P., N.Z., G.C.L., T.H., F.T., and C.L.P.; Visualization, J.L.F., A.S., C.R.S. and C.L.P., Supervision, N.H., G.C.L., T.H., F.T., and C.L.P; Project Administration, F.T. and C.L.P; Funding Acquisition, N.Z., T.H., F.T., and C.L.P.

## Methods

### Protein expression and purification

Using the baculovirus expression system (Invitrogen), His-tagged mouse CRY1, CRY2, CRY2 7M, CLOCK:BMAL1 bHLH-PAS, or GST-tagged BMAL1 PAS-AB were expressed in Sf9 suspension insect cells (Expression Systems). Sf9 suspension cells were infected with a P3 virus at 1.2 × 10^6^ cells per milliliter and grown for 72 hours. Cells were centrifuged at 4°C for 4000 x rpm for 15 minutes. CRY1, CRY2 and CRY2 7M cells were resuspended in 50 mM Tris buffer pH 7.5, 300 mM NaCl, 20 mM imidazole, 10% (vol/vol) glycerol, 0.1% (vol/vol) Triton X-100, 5 mM β-mercaptoethanol and EDTA-free protease inhibitors (Pierce). Cells were lysed using a microfluidizer followed by sonication with a ¼” probe on ice for 15 seconds on, 45 seconds off for three pluses at 40% amplitude. Lysate was clarified at 4°C for 19,000 rpm for 45 minutes. The protein was isolated by Ni2+-nitrilotriacetic acid (Ni-NTA) affinity chromatography (Qiagen). Proteins were eluted with 50 mM Tris buffer pH 7.5, 300 mM NaCl, 250 mM imidazole, 10% (vol/vol) glycerol, and 5 mM β-mercaptoethanol. Proteins were further purified by HiTrap HP cation exchange chromatography (GE Healthcare) and Superdex75 gel filtration chromatography (GE Healthcare) into 20 mM HEPES buffer pH 7.5, 125 mM NaCl, 5% (vol/vol) glycerol, and 2 mM tris(2-carboxyethyl)phosphine (TCEP).

CLOCK:BMAL1 bHLH-PAS cells were resuspended in 20 mM sodium phosphate buffer pH 8, 15 mM imidazole, 10% (vol/vol) glycerol, 0.1% (vol/vol) Triton X-100 and 5 mM β-mercaptoethanol. Cells were lysed, clarified and Ni-NTA affinity purification was performed as described above. The complex was eluted with sodium phosphate buffer pH 8, 250 mM imidazole, 10% (vol/vol) glycerol and 5 mM β-mercaptoethanol. The complex was further purified by HiTrap Heparin HP affinity column (GE Healthcare) after diluting eluant ~5-fold with 20 mM sodium phosphate buffer pH 7.5, 50 mM NaCl, 2 mM Dithiothreitol, and 10% (vol/vol) glycerol and loaded onto the column. After washing with 5 column volumes of the above buffer, the complex was then eluted with a gradient 0-100% of 20 mM sodium phosphate buffer pH 7.5, 2 M NaCl, 2 mM Dithiothreitol, and 10% (vol/vol) glycerol. The complex was further purified by Superdex200 gel filtration chromatography (GE Healthcare) into 20 mM HEPES buffer pH 7.5, 125 mM NaCl, 5% (vol/vol) glycerol, and 2 mM TCEP.

GST-BMAL1 PAS-AB-expressing *E. coli* cells were resuspended in 50 mM HEPES buffer pH 7.5, 300 mM NaCl, 5% (vol/vol) glycerol and 5 mM β-mercaptoethanol. Cells were lysed and clarified as described above. The soluble lysate was bound in batch-mode to Glutathione Sepharose 4B (GE Healthcare), then washed and eluted with 50 mM HEPES buffer pH 7.5, 150 mM NaCl, 5% (vol/vol) glycerol and 5 mM β-mercaptoethanol, 25 mM reduced glutathione. The protein was desalted into HEPES buffer pH 7, 150 mM NaCl, 5% (vol/vol) glycerol and 5 mM β-mercaptoethanol using a HiTrap Desalting column (GE Healthcare) and the GST tag was cleaved with GST-TEV protease overnight at 4°C. The cleaved GST-tag and GST-tagged TEV protease was removed by Glutathione Sepharose 4B (GE Healthcare) and the remaining BMAL1 PAS-AB protein was further purified by Superdex75 gel filtration chromatography (GE Healthcare) into 20 mM HEPES buffer pH 7.5, 125 mM NaCl, 5% (vol/vol) glycerol, and TCEP.

Recombinant baculoviruses coding GST-CRY2 PHR, His-CLOCK bHLH-PAS, and His-BMAL1 FL were co-infected into monolayer HighFive (Invitrogen) insect cells for protein expression. Cells were harvested 48 hours after infection, and lysed in a buffer containing 40 mM HEPES pH 7.5, 300 mM NaCl, 10% (v/v) glycerol, 5 mM β-mercaptoethanol. Lysed cells were clarified by ultracentrifugation, and the soluble lysate was purified over a glutathione affinity column (GE Healthcare). Bound complex was eluted via overnight on-column TEV cleavage to remove the GST-and His-tags. Eluted material was further purified by a HiTrap Q-HP (GE Healthcare) anion exchange column, followed by a Superdex 200 gel filtration column (GE Healthcare) equilibrated in 20 mM HEPES pH 7.5, 300 mM NaCl, 10% (v/v) glycerol, 1 mM DTT.

The CLOCK PAS-AB and PER2 CBD were expressed in Rosetta (DE3) *E. coli*. Protein expression was induced with 0.5 mM isopropyl-β-D-thiogalactopyranoside (IPTG) at an OD_600_ of ~0.8, after which cells were grown for an additional 18 hours at 18° C. Cells were harvested by centrifugation at 4°C for 4000 x rpm for 15 minutes. For CLOCK PAS-AB (wild-type or W362A mutant), the protein was expressed as a fusion protein with the solubilizing tag His-NusA-XL and an N-terminal biotin acceptor peptide (BAP) C-terminal to the TEV site. Cells were resuspended in 50 mM Tris buffer pH 7.5, 300 mM NaCl, 20 mM imidazole, 10% (vol/vol) glycerol and 5 mM β-mercaptoethanol and lysed by microfluidizer. Lysate was clarified at 4°C for 19,000 rpm for 45 minutes. The protein was isolated by Ni-NTA affinity chromatography (Qiagen) and eluted with 50 mM Tris buffer pH 7.5, 300 mM NaCl, 250 mM imidazole, 10% (vol/vol) glycerol, and 5 mM β-mercaptoethanol. The protein was then desalted into 50 mM Tris buffer pH 7.5, 150 mM NaCl, 5% (vol/vol) glycerol and 5 mM β-mercaptoethanol, and the His-NusA-XL tag was cleaved with His-TEV protease overnight at 4°C. The cleaved tag and protease were removed by Ni-NTA affinity chromatography (Qiagen) and CLOCK PAS-AB was further purified by Superdex75 gel filtration chromatography (GE Healthcare) into 20 mM HEPES buffer pH 7.5, 125 mM NaCl, 5% (vol/vol) glycerol, and 2 mM TCEP. PER2 CBD was expressed as a fusion protein with GST and purified as described above.

CRY proteins were stored on ice in 20 mM HEPES buffer pH 7.5, 125 mM NaCl, 5% (vol/vol) glycerol, and 2 mM TCEP for up to 7 days. All other proteins were flash frozen and stored at −70°.

### Biotinylation of CLOCK PAS-AB

100 μM BAP-CLOCK PAS-AB (wild-type or W362A mutant) in 20 mM HEPES buffer pH 7.5, 125 mM NaCl, 5% (vol/vol) glycerol, and 2 mM TCEP was incubated at 4°C overnight with 2 mM ATP, 1 μM GST-BirA and 150 μM biotin. GST-BirA was removed afterwards with Glutathione Sepharose 4B (GE Healthcare) resin and excess biotin was separated from the labeled protein by Superdex75 gel filtration chromatography (GE Healthcare) in 20 mM HEPES buffer pH 7.5, 125 mM NaCl, 5% (vol/vol) glycerol, and 2 mM TCEP.

### Assembling protein complexes

To assemble protein:protein or protein:DNA complexes, purified proteins and annealed mouse *Per2* E-box DNA (GCG CGG TCA CGT TTT CCA CT) ~300 μL of 5 μM complex were mixed at a 1:1 molar ratio and incubated for 30 minutes at 4° before injection onto a Superdex200 10/300 GL analytical scale column (GE Healthcare). For CRY:PER2 CBD complexes used for BLI or CRY1:PER2 CBD:CLOCK:BMAL1 bHLH-PAS:Ebox DNA for cryo-EM, a slight molar excess of zinc chloride was added to the protein complexes. The assembly and purity of complexes from peak fractions of the Superdex200 10/300 GL gel filtration column (GE Healthcare) was assessed by SDS-PAGE gel electrophoresis and SimplyBlue SafeStain (Invitrogen) staining.

### Fluorescence polarization

A peptide of the minimal BMAL1 TAD (residues 594-626) was purchased from Bio-Synthesis with a 5,6-TAMRA fluorescent probe covalently attached to the N terminus. Binding assays with CRY1 PHR, CRY1 PHR:PER2 CBD, CRY2 PHR and CRY2 PHR:PER2 CBD were performed in 50 mM Bis-Tris Propane buffer pH 7.5, 100 mM NaCl, 2 mM TCEP and 0.05% (vol/vol) Tween-20. The BMAL TAD probe was diluted to a working concentration of 50 nM in assay buffer and binding was monitored by changes in fluorescence polarization with an EnVision 2103 multilabel plate reader (Perkin Elmer) with excitation at 531 nm and emission at 595 nm. Equilibrium binding dissociation constants (*K*_d_) were calculated by fitting the dose-dependent change in millipolarization (mP) to a one-site specific total binding model in Prism 7 (GraphPad). The mP values shown represent the average of duplicate samples from a representative experiment of n = 3 independent assays. The *K*_d_ reported (± sd) is the average of determined from the 3 independent assays.

### Bio-layer interferometry

All BLI experiments were performed using an 8-channel Octect-RED96e (ForteBio) with a BLI assay buffer of 20 mM HEPES buffer pH 7.5, 125 mM NaCl, 5% (vol/vol) glycerol and 2 mM TCEP. All experiments began with a reference measurement to establish a baseline in BLI buffer for 120 seconds. Next, 1.5 µg/mL biotinylated CLOCK:BMAL1 PAS-AB (wild-type or W362A mutant in BLI buffer) was loaded on a streptavidin tip for 300 seconds. Subsequently, a 360 second blocking step was performed with 0.5 mg/mL BSA, 0.02% (vol/vol) Tween, 20 mM HEPES buffer pH 7.5, 125 mM NaCl, 5% (vol/vol) glycerol and 2 mM TCEP. Association was then measured for 300 seconds for 8 different concentrations of the analyte (CRY, CRY:PER2 CBD, PER2 CBD) in a serial dilution starting at approximately 10x the estimated *K*_d_ in blocking buffer. Dissociation was measured for 300 seconds in blocking buffer. Each experiment was repeated with tips that were not loaded with CLOCK:BMAL PAS-AB to provide a reference for non-specific binding to the tip. Data were processed and fit using Octet software v.7 (ForteBio). Before fitting, all datasets were reference-subtracted, aligned on the y-axis and aligned for interstep correction through their respective dissociation steps according to the manufacturer’s instructions. For each experiment, at least 4 different concentrations were used to fit association and dissociation globally using a 1:1 binding model in Octet software v.7 (ForteBio). Ultimately, the goodness of fit was determined using χ^2^ and R^2^ values according to the manufacturer’s instructions.

### Molecular dynamics simulations

The crystal structure of apo CRY1 (PDB: 5T5X) and apo CRY2 (PDB: 4I6E) were prepared for starting models for the molecular dynamics (MD) simulations by modeling non-terminal missing residues using the Prime program (Schrödinger). All crystallographically defined water molecules were removed from the structures. In order to prepare the system with the *in silico* mutated CRY2 (A61G/S64N CRY2), the corresponding amino acids were mutated in wild type apo CRY2 using UCSF-Chimera (Pettersen et al., 2004) and used as starting models. Amber99sb-ildn force field was used for simulation (Lindorff-Larsen et al., 2010). The structures were solvated in a dodecahedron box using TIP3P water molecules. The systems were neutralized and then Na^+^ and Cl^−^ ions were added to maintain a physiological ionic concentration of 0.15 M. Energy minimization was performed for the systems until the maximum force on any atom was less than 1,000 kJ mol^−1^nm^−1^ in the case of apo CRY1 and CRY2, and 500 kJ mol^−1^nm^−1^ in the case of the *in silico* mutated CRY2.

After energy minimization, the systems were equilibrated in an NVT ensemble for 500 ps. The temperature was maintained at 310 K using the modified Berendsen Thermostat (V-rescale) (Bussi et al., 2007). After this, the systems were equilibrated in an NPT ensemble for 500 ps. The pressure was maintained at 1 bar using a Berendsen barostat (Berendsen et al., 1984). This was followed by a production run of 500 ns for each system in an NPT ensemble using a Parinello-Rahman barostat (Parrinello and Rahman, 1981) and modified Berendsen Thermostat (V-rescale) with the temperature maintained at 310 K and pressure at 1 bar. Three independent simulations were completed for the apo CRY1 and apo CRY2 systems described in the previous section and two independent simulations were run for the *in silico* mutated CRY2 system. All MD simulations were performed using Gromacs 5 (Abraham et al., 2015).

The trajectories were analyzed using in built functions of Gromacs. RMSD, RMSF, distances and dihedral angles were calculated using gmx rmsd, gmx rmsf, gmx distance and gmx chi programs of Gromacs respectively. UCSF-Chimera (Pettersen et al., 2004) was used for molecular visualization. The volume of secondary pocket for 251 equally spaced trajectory frames from each run was calculated using POVME 3.0 (Wagner et al., 2017). The comparison of volumes between CRY1-CRY2 and between CRY2-Mut CRY2 was done by performing Wilcoxon ranksum test as implemented in Scipy (https://www.scipy.org/).

### Calculation of Kullback-Leibler (KL) divergence

The KL-divergence or relative entropy has been previously used to quantify the residue wise differences between the ensembles (McClendon et al., 2012; Moffett et al., 2017; Rapp et al., 2013). To calculate the KL-divergence between the ensembles using mutinf software (McClendon et al., 2012), we divided each simulation trajectory between 100 ns and 500 ns into 4 blocks of 100 ns each. This gave a total of 12 blocks for apo CRY1 and apo CRY2 simulations and 8 blocks for the *in silico* mutated CRY2. Dihedral angles phi, psi and chi1 were calculated for 5000 equally spaced frames from each block and used as input to calculate KL-divergence. Dividing the trajectories into multiple blocks of sufficiently large number of frames enabled the calculation of bootstrap values, which were used to make a robust estimation of significant differences between the two ensembles. The divergence values greater than the bootstrap values for a given residue suggests that the dynamics of those residues differ significantly between the two ensembles, in terms of side chain rotamer distribution, main chain dihedral angle distribution or both.

### Crystallization, data collection and structure determination

The CRY1 PHR:PER2 CBD complex was purified as described above. The protein was concentrated to 4.3 mg/mL and crystalized by sitting-drop vapor diffusion at 22°C. Crystals formed in a 1:1 ratio of protein to precipitant in 0.2 M MgCl_2_, 0.1 M Bis-Tris buffer pH 5.5 and 25% (vol/vol) PEG 3350. Crystals were harvested and flash frozen in the reservoir solution with 20% (vol/vol) glycerol before data collection. Data were collected from single crystal at λ = 1.0A, 100 K on Beamline 23-ID-D at the Advanced Photon Source (Argonne, Illinois, USA). Diffraction images were indexed and scaled using iMosflm (Battye et al., 2011) and Scala (Evans, 2006) in the CCP4 package (Winn et al., 2011). Phases were solved by molecular replacement with PHASER (Adams et al., 2010) using the crystal structure of mouse CRY1 in complex with the PER2 CBD (PDB: 4CT0). All reflections were used for refinement except for 5% that were excluded for R_free_ calculations. The structural model was revised in real space with the program COOT (Emsley et al., 2010) based on sigma-A weighted 2Fo-Fc and Fo-Fc electron density maps. Data collection and final refinement statistics are given in Table S1.

### Sample preparation for electron microscopy

The full complex containing CLOCK:BMAL1 bHLH-PAS bound to an annealed DNA duplex containing the minimal mouse *Per2* E-box (GCG CGG TCA CGT TTT CCA CT), and CRY1 PHR:PER2 CBD was briefly incubated on ice in 20 mM Hepes, 125 mM NaCl, 2 mM TCEP and 1% (vol/vol) glycerol. 2.5 µL of sample was then applied to an UltraAuFoil R1.3/1.3 300-mesh grid (Electron Microscopy Services), which was freshly plasma-cleaned using a Gatan Solarus (75% argon/2% oxygen atmosphere, 15 W for 7 seconds). Grids with applied sample were then manually blotted with filter paper (Whatman No.1) for ~3 seconds in a 4°C cold room before plunge freezing in liquid ethane cooled by liquid nitrogen.

### Electron microscopy data acquisition

The Leginon automated data-acquisition program (Suloway et al., 2005) was used to acquire all cryo-EM data. Real-time image pre-processing, consisting of frame alignment, contrast transfer function (CTF) estimation and particle picking, were performed using the Appion image-processing pipeline during data collection (Voss et al., 2009). Image collection was performed using a Thermo Fischer Talos Arctica operating at 200 keV and equipped with a Gatan K2 Summit DED, at a nominal magnification of 36,000X (corresponding to a physical pixel size of 1.15 Å per pixel). 692 movies were collected, with 48 frames per movie using a total exposure time of 12 seconds with an exposure rate of 5.2 e^−^/pixel/s and total exposure of 47.2 e^−^/Å^2^ (0.98 e^−^ per frame). A nominal defocus range from −0.8 μm to −1.8 μm was used during data collection.

### Electron microscopy image processing

Automated particle picking was conducted using the Difference of Gaussians (DoG) picker (Voss et al., 2009) to yield 891,700 particle picks. Before particle extraction, CTFFIND4 was used for CTF estimation. Fourier-binned 2×2 particles were subjected to reference-free two-dimensional (2D) classification in Relion 3.0 (Scheres, 2012) to remove poorly aligning particles and non-particles in the data. A second round of 2D classification in Relion was used to further sort out bad particles and false-picks, resulting in 81,257 particles populating 2D classes with strong structural features to be used for further processing (Supplementary Fig. 4a). This subset of particles was then subjected to reference-free ab initio model generation in cryoSPARC (Punjani et al., 2017) which then underwent one round of heterogenous refinement into three classes. One class appeared more ordered compared to the other two and consisted of the largest proportion of particles (44%). This map was rescaled to 1.15 Å per pixel and was used as an initial model for Relion 3D classification using the extracted and unbinned original subset of particles selected from the two rounds of 2D classification, resulting in three classes. The class with the largest proportion of particles (50% or 40,628 particles) appeared superior from visual inspection and underwent one round of 3D autorefinement in Relion resulting in a 7.2 Å map. This was followed by one round of CTF Refinement and postprocessing in Relion resulting in a final map with a reported global resolution of ~6.1 Å. Some anisotropy was observed in our cryo-EM density, arising from particles adopting two preferred orientations in vitrified ice (Figure S6), which was detrimental to the resolution of our 3D reconstruction and likely leads to an inflated value for our final map. This is likely arising from interaction with the air-water interface with our complex, as commonly seen in other protein samples analyzed by cryo-EM (Glaeser, 2016; Glaeser and Han, 2017; Noble et al., 2018). Attempts to improve the particle distribution by addition of detergents to the sample or by stage-tilting during data collection (Tan et al., 2017) were ineffective and did not result in a higher quality map. Despite this, some secondary structural features were resolvable, providing features for docking of crystal structures and the eventual generation of a pseudoatomic model.

### HADDOCK studies

The cluster representatives obtained after clustering the trajectories were used to dock the PAS-B domain (residues 262-384) of CLOCK (PDB: 4F3L) using HADDOCK 2.2 webserver (van Zundert et al., 2016). The active residues used for generating restraints for docking were: D38/D56, P39/P57, F41/F59, R51/R69, G106/G124, R109/R127, F257/F275, E382/D400, E383/E401 for CRY1/CRY2 and G332, H360, Q361, W362, E367 for CLOCK. The passive residues were specified as residues surrounding the active residues. The PAS-B domain of the CLOCK protein was used to dock to ensemble of structures comprising of 15 cluster representatives of apo CRY1 and 12 cluster representatives of apo CRY2. A total of 10000, 400 or 400 models were generated, respectively, for the rigid docking, flexible docking and water refinement phase of the docking protocol. The final 400 models were clustered using Fractional Common Contacts (FCC) and sixteen clusters were obtained comprising 360 structures. Out of these sixteen, there were three large clusters with 92, 80 and 60 complexes constituting about 65% of the clustered structures. The representatives from these three largest clusters were considered for further analysis.

### Flexible fitting of CRY1 PHR:PER2 CBD:CLOCK PAS-B complex to EM map

The complex comprising CRY1 PHR:PER2 CBD (PDB: 6OF7) and CLOCK PAS-B from a representative structure of the HADDOCK Cluster 1 was fit into the EM map as rigid body using Situs (Wriggers, 2012). The best fitting model was then used to perform flexible fitting. Briefly, the rigid fit model was first converted to an all-atom structure based model (SBM) using SMOG server (Noel et al., 2016). Molecular dynamics simulation was then performed on this all-atom SBM model with a biasing potential based on the EM Map (Orzechowski and Tama, 2008) using MDFit (Whitford et al., 2011) packaged in gromacs 4.5.5 (Pronk et al., 2013).

### Software References

SMOG webtool – http://smog-server.org/

MDFit – http://smog-server.org/extension/MDfit.html

Gromacs – http://www.gromacs.org/

