## Supplemental Figures and Tables for "Protein dynamics regulate distinct biochemical properties of cryptochromes in mammalian circadian rhythms"

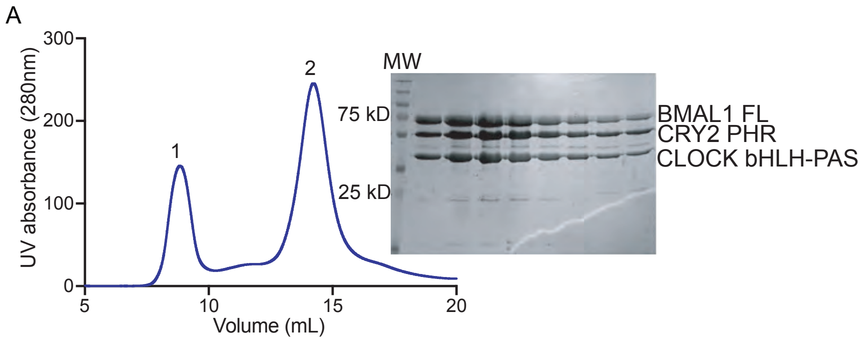


Figure S1, related to Figure 2 – Gel filtration analysis of CRY1, CRY2 and CLOCK:BMAL bHLH-PAS proteins.
A, Chromatographic analysis of complex formation by CRY2 PHR and a heterodimer of CLOCK bHLH-PAS with full-length BMAL1, as above. The addition of the BMAL1 C-terminus resulted in co-elution of CLOCK:BMAL1 with CRY2 PHR in peak 2 (peak 1 contains aggregated protein eluting in the void volume).

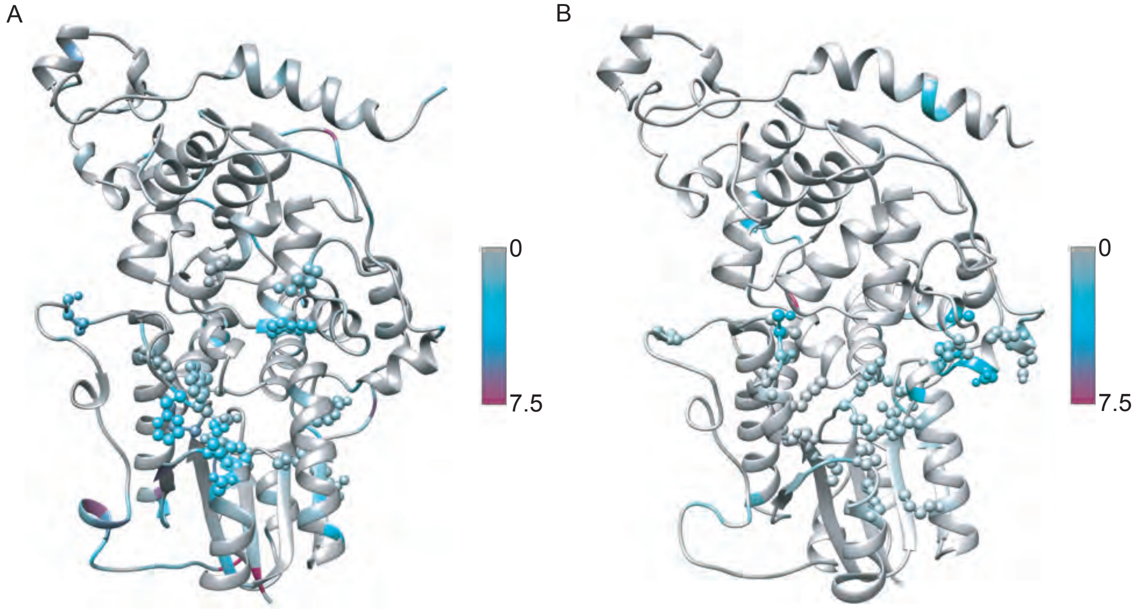


Figure S2, related to Figure 3 – Several residues show high KL divergence between CRY1 and CRY2. A, KL divergence between CRY1 and CRY2 ensembles. The residues are colored as per the KL divergence. The color bar shows values of the KL divergence corresponding to different colors. B, KL divergence between CRY2 and Mut-CRY2 ensembles. Residues showing high KL divergence near the secondary pocket are displayed in ball and stick representation.

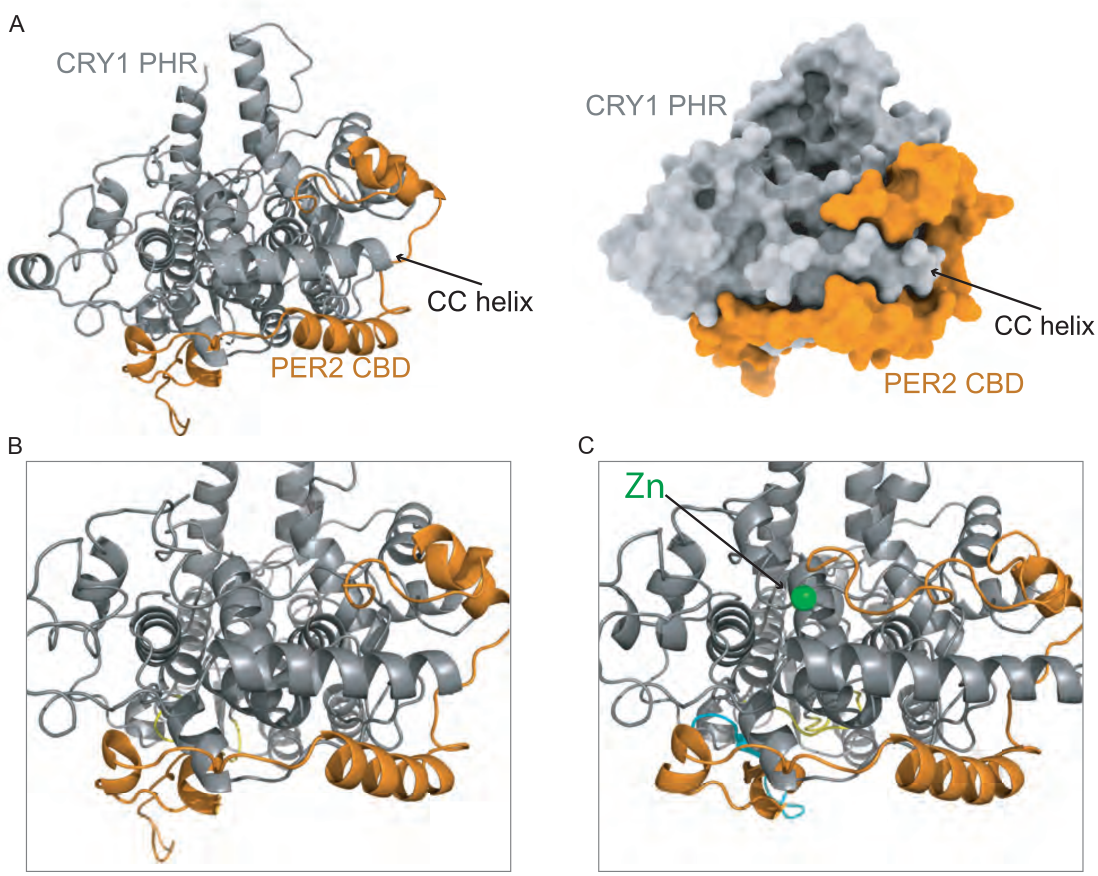


Figure S3, related to Figure 4 – CC helix view of the new CRY1 PHR:PER2 CBD crystal structure. A, Crystal structure of CRY1 PHR:PER2 CBD. CRY1 PHR is shown in gray and PER2 CBD is shown in orange. Ribbon (left) and surface (right) representations of the CC helix, where the TAD of BMAL1 binds. B, Zoomed in view of the CC helix. C, Zoomed in view of the CC helix of the previously solved crystal structure of CRY1 PHR:PER2 CBD (4TC0). CRY1 PHR is shown in gray and PER2 CBD is shown in orange. The far C-terminus of the CBD of PER is coordinating a zinc ion (green) with CRY1 PHR. The zinc ion and subsequent coordinating residues are missing from the new structure.

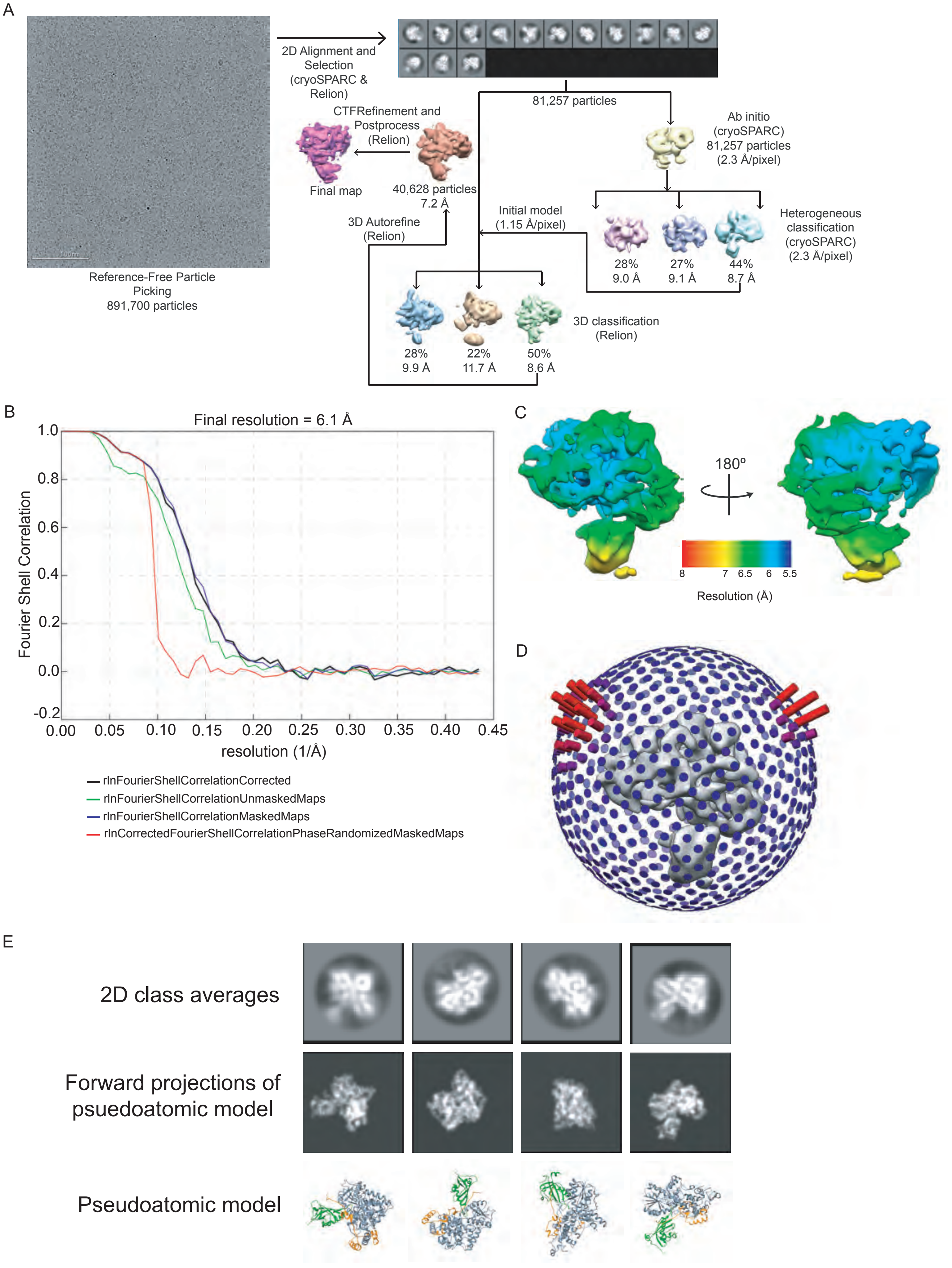
A
2D Alignment and Selection (cryoSPARC & Relion)
Reference-Free Particle Picking
891,700 particles
81,257 particles
Ab initio (cryoSPARC) 81,257 particles (2.3 Å/pixel)
CTFRefinement and Postprocess (Relion)
Final map
40,628 particles
7.2 Å
3D Autorefine (Relion)
Initial model (1.15 Å/pixel)
Heterogeneous classification (cryoSPARC) (2.3 Å/pixel)
28%
9.0 Å
27%
9.1 Å
44%
8.7 Å
3D classification (Relion)
28%
9.9 Å
22%
11.7 Å
50%
8.6 Å
B
Final resolution = 6.1 Å
Fourier Shell Correlation
1.0
0.8
0.6
0.4
0.2
0.0
-0.2
0.00
0.05
0.10
0.15
0.20
0.25
0.30
0.35
0.40
0.45
resolution (1/Å)
rlnFourierShellCorrelationCorrected
rlnFourierShellCorrelationUnmaskedMaps
rlnFourierShellCorrelationMaskedMaps
rlnCorrectedFourierShellCorrelationPhaseRandomizedMaskedMaps
C
180º
8
7
6.5
6
5.5
Resolution (Å)
D
E
2D class averages
Forward projections of psuedoatomic model
Pseudoatomic model

Figure S4, related to Figure 5 – Image processing workflow, global and local resolution of the final map and model validation.
A, Image processing workflow for single particle cryo-EM data. B, Gold standard FSC plot generated for final sharpened map. C, Local resolution plot for final sharpened map. D, Euler plot showcasing particle distribution present in final subset of particles contributing to the final map. Less populated views are shown in blue, while more populated views are depicted in red. E, Comparison between 2D class averages (top row) from the cryo-EM data and our pseudoatomic model of CRY PHR:PER2 CBD bound to CLOCK PAS-B. Forward projections (middle row) of the pseudoatomic model (bottom row) were generated to match the angular orientations observed in the experimental data.

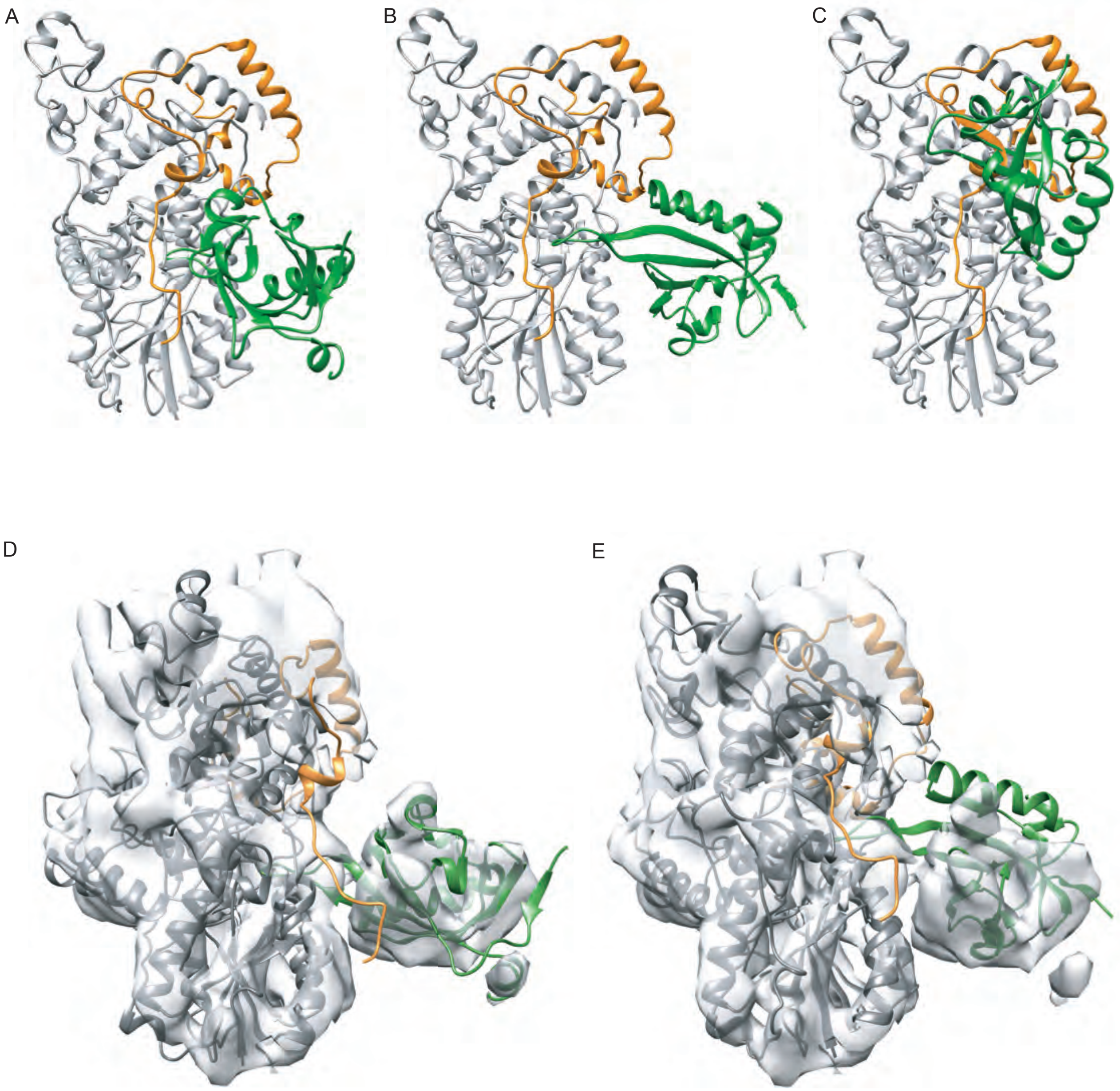


Figure S5, related to Figure 5 - Superimposition of CRY1 PHR:PER2 CBD on the representative models from CRY1:CLOCK PAS-B haddock docking clusters and rigid fit models of CRY1 PHR:PER2 CBD:CLOCK PAS-B on EM map.

A, Cluster 1 representative in complex with PER2 CBD. B, Cluster 2 representative in complex with PER2 CBD. C, Cluster 3 representative in complex with PER2 CBD. D, The model obtained by fitting Cluster 1. E, The model obtained by fitting Cluster 2.

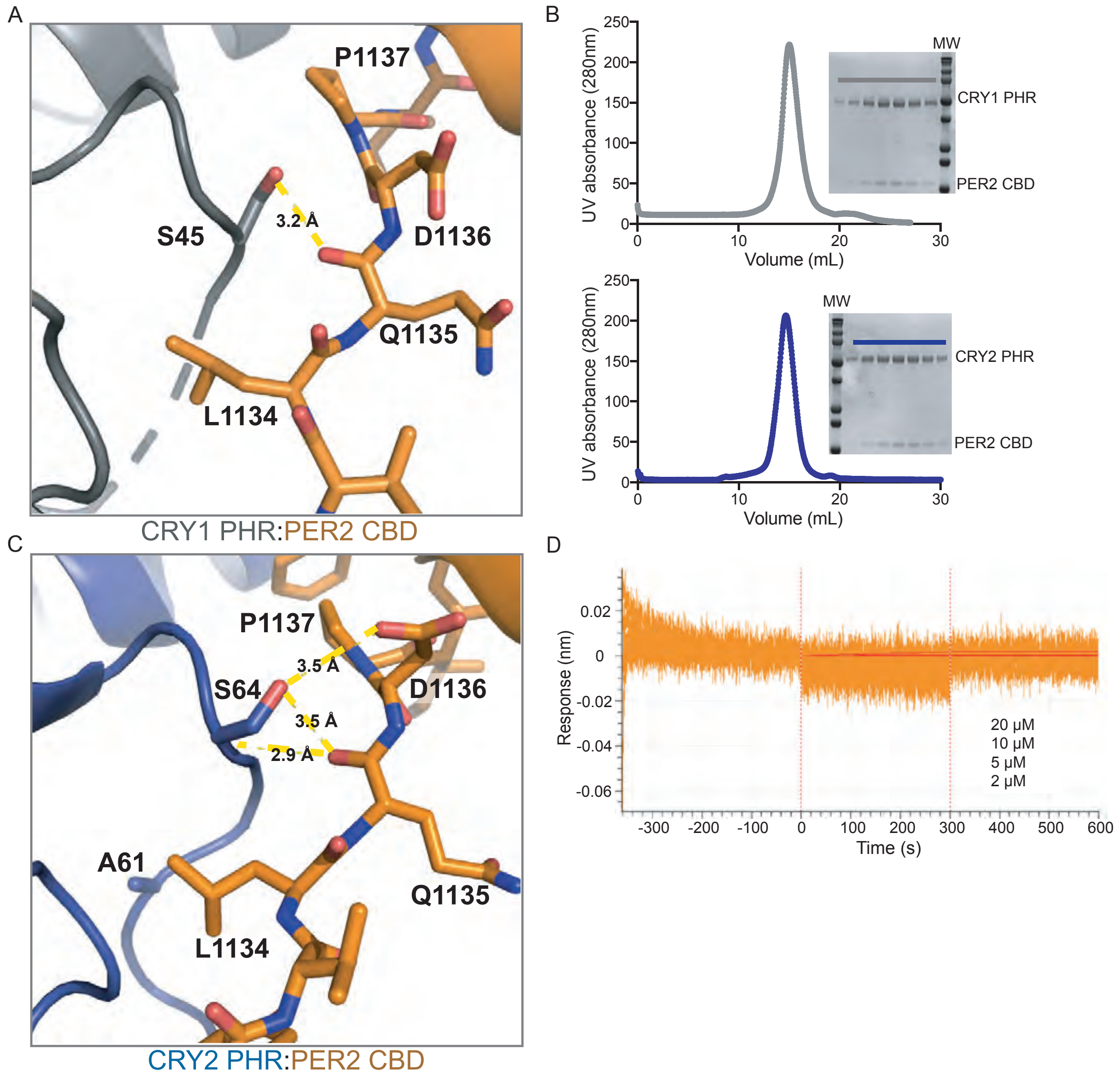


Figure S6, related to Figure 6 – Dimerization with PER2 CBD rearranges the serine loops on CRY1 and CRY2.
A, Zoomed in view of the serine loop of CRY1 PHR:PER2 CBD (gray:orange). Interaction between Ser45 of CRY1 and the backbone of Gln1135 of PER2 CBD shown with yellow dashed lines. B, Gel filtration analysis of CRY1 PHR:PER2 CBD (top, gray) and CRY2 PHR:PER2 CBD (bottom, blue). Protein was run on a S200 10/300 GL column and peak fractions were analyzed by SDS/PAGE. C, Zoomed in view of the serine loop of CRY2 PHR:PER2 CBD (blue:orange). Interaction between Ser64 of CRY2 and the backbone of Gln1135 and the side chain of Asp1136 of PER2 CBD shown with yellow dashed lines. D, Processed bio-layer interferometry (BLI) data for PER2 CBD (orange) binding to biotin labeled CLOCK:BMAL PAS-AB. Inset values represent the concentrations of CRY for the individual binding reactions, top to bottom. The red line is the nonlinear least squares fitting. PER2 CBD does not bind to CLOCK:BMAL1 PAS-AB at the concentrations tested.

### Supplemental Tables

Supplemental Table 1. X-ray crystallography data collection and refinement statistics. Values in parentheses are for highest resolution shell. Related to Figure 4.

| | **CRY1 PHR:PER2 CBD** |
|---|---|
| **Data collection** | |
| Space group | P $2_1$ $2_1$ $2_1$ |
| Cell dimensions | |
| a, b, c | 72.64, 81.23, 98.10 |
| $\alpha$, $\beta$, $\gamma$ | 90, 90, 90 |
| Resolution (Å) | 62.56 – 3.11 (3.28-3.11) |
| $R_{merge}$ | 17.1 (65.2) |
| Total reflections | 62925 (9485) |
| Unique reflections | 10939 (1564) |
| $CC_{1/2}$ | 0.99 (0.75) |
| $I/\sigma$ | 7.5 (3.0) |
| Completeness | 100 (100) |
| Redundancy | 5.8 (6.1) |
| **Refinement** | |
| $R_{work}/R_{free}$ | 20.8/26.2 |
| Number of atoms | 4424 |
| Protein | 4424 |
| Water | - |
| Ligands | - |
| Bond lengths | 0.003 |
| Bond angles | 0.43 |
| Average B factor | 60.72 |
| Ramachandran Favored/Outliers (%) | 99.45/0.55 |

Supplemental Table 2. Haddock docking details related to Figure 5.

| **Cluster ID** | **1** | **2** | **3** |
|---|---|---|---|
| Cluster Size | 92 | 80 | 60 |
| z-score | -0.4 | -1.9 | -1.4 |
| RMSD from lowest energy structure | 11.7 ± 0.3 | 0.8 ± 0.5 | 14.5 ± 0.4 |
| Van der Waals | -64.5 ± 4.2 | -73.5 ± 4 | -70.9 ± 7.8 |
| Electrostatic | -241.6 ± 29.2 | -230.8 ± 23.1 | -177.9 ± 21.0 |
| Desolvation | -22.9 ± 6.0 | -45.3 ± 4.8 | -48.6 ± 3.9 |
| Restrains Violation | 3.7 ± 2.0 | 52 ± 20.4 | 31.1 ± 25.7 |
| Buried Surface Area | 1841.9 ± 61.8 | 2134.9 ± 106.1 | 2126.0 ± 98.7 |
